# Structural basis for the RNA-guided ribonuclease activity of CRISPR-Cas13d

**DOI:** 10.1101/314401

**Authors:** Cheng Zhang, Silvana Konermann, Nicholas J. Brideau, Peter Lotfy, Scott J. Novick, Timothy Strutzenberg, Patrick R. Griffin, Patrick D. Hsu, Dmitry Lyumkis

## Abstract

CRISPR-Cas endonucleases directed against foreign nucleic acids mediate prokaryotic adaptive immunity and have been tailored for broad genetic engineering applications. Type VI-D CRISPR systems contain the smallest known family of single effector Cas enzymes, and their signature Cas13d ribonuclease employs guide RNAs to cleave matching target RNAs. To understand the molecular basis for Cas13d function, we resolved cryo-electron microscopy structures of Cas13d-guide RNA binary complex and Cas13d-guide-target RNA ternary complex to 3.4 and 3.3 Å resolution, respectively. Furthermore, a 6.5 Å reconstruction of apo Cas13d combined with hydrogen-deuterium exchange revealed conformational dynamics that have implications for RNA scanning. These structures, together with biochemical and cellular characterization, explain the compact molecular architecture of Cas13d and provide insights into the structural transitions required for enzyme activation. Our comprehensive analysis of Cas13d in diverse enzymatic states facilitated site-specific truncations for minimal size and delineates a blueprint for improving biomolecular applications of RNA targeting.

## Introduction

Bacterial life employs diverse immune mechanisms to protect themselves against predatory phage, which are thought to outnumber them by ten to one (Chibani-Chennoufi et al., 2004). These include abortive infection (Abi), toxin-antitoxin (T-AT) modules, and direct cleavage of phage nucleic acids using restriction enzymes and CRISPR-Cas systems. CRISPR systems in particular engage their constituent Cas nucleases with programmable guide RNAs to target invading nucleic acids, endowing the host cell with adaptive immunity (Barrangou et al., 2007; Brouns et al., 2008; Gasiunas et al., 2012; Jinek et al., 2012). They are broadly divided into two classes, each with multiple types and subtypes, wherein Class 1 CRISPR systems (types I and III) coordinate multiple proteins that cooperate for target surveillance and defense, while Class 2 systems integrate both functions into a single effector enzyme (Hille et al., 2018; Makarova et al., 2015).

Class 2 CRISPR-Cas systems include types II, V, and VI, with types II and V shown to target DNA (Shmakov et al., 2017). Adapted over the last half-decade into a remarkably flexible genetic engineering toolbox, Class 2 DNA-targeting enzymes such as CRISPR-Cas9 (type II) and CRISPR-Cas12a/Cpf1 (type V) have facilitated many applications, from gene editing to lineage tracing, multi-color chromosomal imaging, and gene drives. Although some Class 1 CRISPR systems can target RNA (Hale et al., 2009; Jiang et al., 2016; Kazlauskiene et al., 2017; Niewoehner et al., 2017; Niewoehner and Jinek, 2016; Samai et al., 2015), Type VI systems have been recently described as the only known single effector RNA-targeting CRISPR nucleases (Abudayyeh et al., 2016; East-Seletsky et al., 2016; Konermann et al., 2018; Shmakov et al., 2015; Smargon et al., 2017; Yan et al., 2018). Cas13, the signature single-effector enzyme family, comprises guide RNA-directed ribonucleases with 4 subtypes (Cas13a-d) that each exhibit significant sequence divergence apart from two consensus HEPN (Higher eukaryotes and prokaryotes nucleotide-binding domain) RNase motifs, R-X_4-6_-H. Domains belonging to the HEPN superfamily are frequently found in ribonucleases involved in immune defense (Anantharaman et al., 2013), including in Class 1 CRISPR RNases such as Csm6 or the homologous Csx1 (Jiang et al., 2016; Kazlauskiene et al., 2017; Niewoehner and Jinek, 2016), as well as prokaryotic Abi and T-AT defense systems or the anti-viral mammalian RNase L (Han et al., 2014). To defend against viral infection, Cas13 enzymes employ two distinct RNase activities to process pre-crRNA into mature crRNA guides in a HEPN-independent manner, followed by HEPN-dependent cleavage of a complementary “activator” target RNA in *cis*. Upon target-dependent activation, Cas13 is also able to cleave bystander RNAs in vitro, reflecting a general RNase activity capable of both *cis*- and *trans*-cleavage.

## Diversity of Cas13 ribonucleases

Despite functional similarities of crRNA-dependent activation of HEPN-mediated RNA cleavage, Cas13 subtypes characterized to date exhibit key differences beyond their significant divergence at the primary sequence level. Using a computational pipeline for identifying novel Class 2 CRISPR-Cas loci from genome and metagenome sequences sourced from large-scale microbiome sequencing efforts, we recently described the discovery of a new Cas13 subtype designated as Cas13d (Konermann et al., 2018). Cas13d enzymes are prominently smaller than other Cas13 subtypes, at an average size of 930 aa compared to 1120-1250 aa for other Cas13s, facilitating flexible packaging into size-constrained therapeutic viral vectors such as adeno-associated virus (AAV) (Konermann et al., 2018; Yan et al., 2018).

Furthermore, while Cas13a and Cas13b exhibit variable dependence on a protospacer-flanking sequence (PFS) for efficient targeting, Cas13d enzymes lack PFS requirements (Konermann et al., 2018; Yan et al., 2018). Some Cas13b and Cas13d systems also associate with regulatory proteins that modulate their RNase activity (Smargon et al., 2017; Yan et al., 2018). Like all CRISPR-Cas effectors described to date, Cas13 enzymes each recognize a direct repeat (DR) sequence containing a conserved stem loop structure within their cognate crRNA. However, between Cas13 subtypes, the DR sequence motifs, RNA fold, and DR position relative to the spacer sequence are each distinct. For Cas13a and Cas13d, the DR is located on the 5′ end (Abudayyeh et al., 2016; Konermann et al., 2018; Yan et al., 2018) while the Cas13b DR is 3′ of the spacer sequence (Smargon et al., 2017).

Cas13 enzymes provide a rich resource for new RNA targeting technologies, and have been recently developed for RNA knockdown (Abudayyeh et al., 2016; Cox et al., 2017; Konermann et al., 2018), editing (Cox et al., 2017), splicing, and viral delivery (Konermann et al., 2018). Remarkably, Cas13 subtypes and individual orthologs exhibit highly variable activity in human cells, with Cas13d overall displaying the most robust reported activity for both target cleavage and binding (Cox et al., 2017; Konermann et al., 2018).

Here, we sought to understand the molecular and structural basis for Cas13d function, including both guide and target RNA recognition. We report cryo-EM reconstructions of 3 distinct structures: *Eubacterium siraeum* Cas13d (*Es*Cas13d) bound to guide RNA at 3.4 Å resolution (binary), *Es*Cas13d in complex with guide RNA and target RNA at 3.3 Å resolution (ternary), and a reconstruction for the stable region of *Es*Cas13d alone (apo). Our structural and biochemical data reveal insights into the function of CRISPR/Cas effector enzymes, including how Cas13 RNA targeting properties are compacted into the minimal Cas13d subtype, and provide a comprehensive blueprint for the rational structure-based modification of Cas13d orthologs for transcriptome engineering.

## Results

### Determination of a high-resolution cryo-EM structure of Cas13d in complex with crRNA

To gain structural insight into Cas13d function, we purified the catalytically active *Es*Cas13d (Konermann et al., 2018) and formed a binary complex containing Cas13d bound to CRISPR RNA (crRNA) followed by cryo-EM imaging (**Figure S1A-B**). A large data collection, followed by a computational analysis and refinement of 48,795 particles led to the derivation of a coulombic potential map describing the binary complex bound to crRNA, resolved to a mostly homogeneous resolution of ~3.4 Å (**Figure S1C-D, and Supplementary Table 1**). Using recently described procedures for characterizing anisotropy in cryo-EM density maps (Tan et al., 2017), we found that the structure maintained approximately even distribution of directional resolution (**Figure S1E-G**).

At 954 amino acids (molecular weight ~105 kDa), *Es*Cas13d is considerably smaller than average members of the Type VI-A, −B, and −C subtypes (Konermann et al., 2018; Yan et al., 2018). Most residues could be built into the density, with the exception of several flexible loops. All 52 nucleotides spanning the crRNA were observed in the EM density and 51 of them could be confidently modeled. The final model is consistent with the cryo-EM map, with good geometry and model statistics (**Table S1**).

### The structure of crRNA-bound Cas13d reveals a compact protein architecture surrounding solvent-exposed RNA

The single effector Cas13d binary complex (**Figure 1A-D**) maintains a bilobed architecture with five distinct domains organized around the central crRNA guide (**Figure 1A**). The domains include an N-terminal domain (NTD), a HEPN1 catalytic domain that is split into two distinct regions in sequence space, the first linker domain termed Helical-1, a second HEPN2 catalytic domain, and a second linker domain termed Helical-2. With the exception of the NTD, which is composed of two short α-helices flanking a β-sandwich region formed by two antiparallel 3-stranded β-sheets, the protein is predominantly alpha-helical.

**Figure 1.**
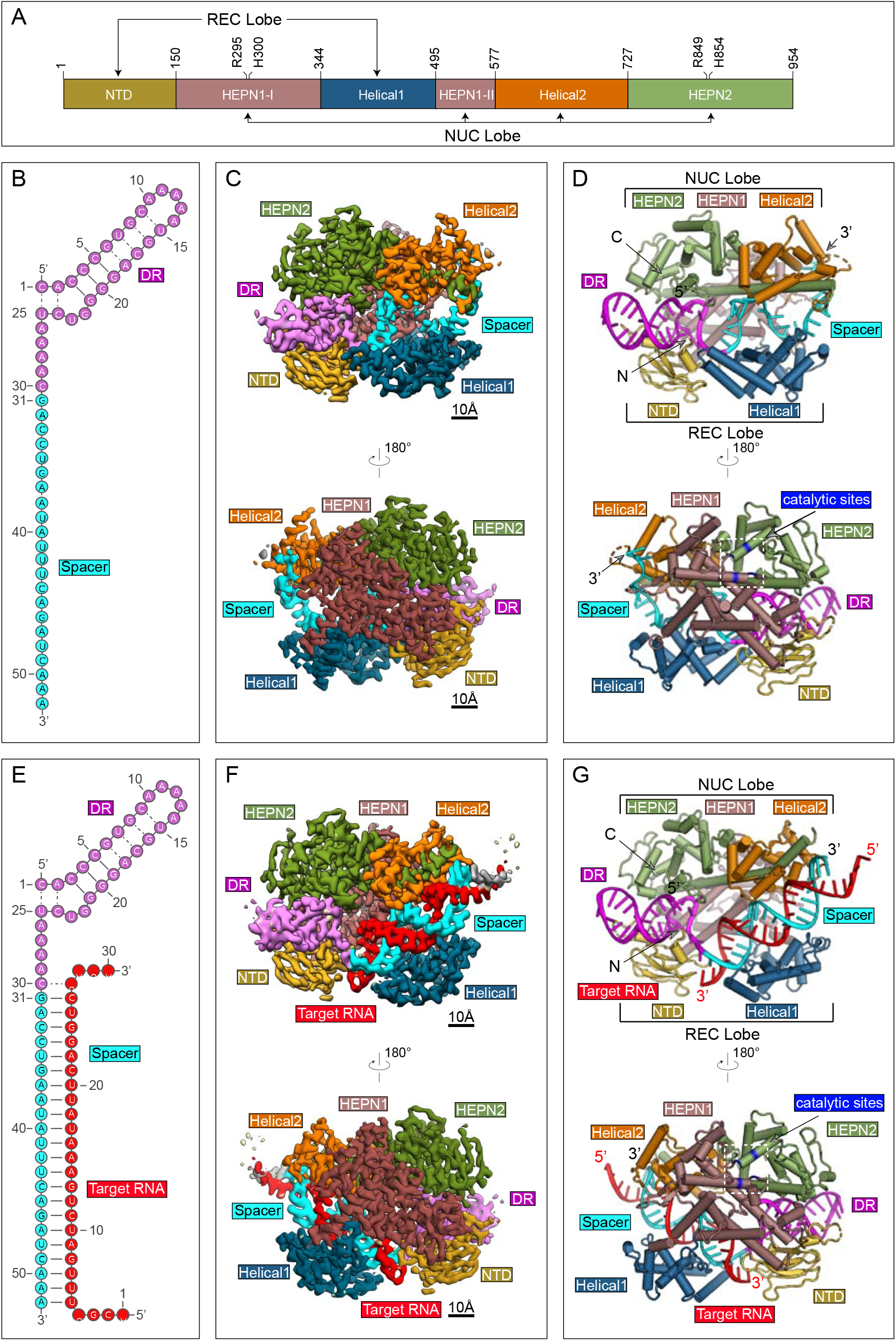
Structures of Cas13d-crRNA and Cas13d-crRNA-target RNA. (A) Domain organization of *Es*Cas13d, with boundaries and catalytic sites of the HEPN domains (R295, H300, R849, H854) indicated. (B) Schematic representation of the crRNA used for binary complex assembly. The DR (5′ handle) and spacer of crRNA are colored magenta and cyan, respectively. (C) Opposing views of the cryo-EM binary reconstruction, resolved to ~3.4 Å resolution. (D) Opposing views of the binary atomic model derived from the cryo-EM density. (E) Schematic representation of the crRNA used for ternary complex assembly. The DR and spacer of crRNA are colored as in (A), target RNA is red. (F) Opposing views of the cryo-EM ternary reconstruction, resolved to ~3.3 Å resolution. (G) Opposing views of the ternary atomic model derived from the cryo-EM density.

The overall binary ribonucleoprotein architecture is reminiscent of a half-open clam shape surrounding the solvent exposed crRNA channel. The mature crRNA is divided into a constant direct repeat (DR) region (nt 1-30), derived from the characteristic repeat of CRISPR arrays, and a spacer region (nt 31-52) complementary to the target protospacers (**Figure 1B**). In the binary complex, the 5′ crRNA handle (also referred to as the DR) is clamped by NTD and HEPN2, with the first two base pairs and its 5-nt loop protruding away from protein density **(Figure 1C, 1D)**. Immediately downstream of the DR, the spacer region resides within a cleft and is sandwiched between Helical-1 and Helical-2. HEPN1 provides a structural scaffold connecting the two lobes of Cas13d, reminiscent of a hinge around the largely solvent-exposed RNA density. In this compact configuration, Cas13d forms a “surveillance complex”, poised for searching and identifying complementary target sites (**Figure 1D**).

### Determination of a high-resolution cryo-EM structure of Cas13d in complex with crRNA and target RNA

Type VI CRISPR-Cas RNases catalyze degradation of ssRNA through a process that is mediated by the formation of an activated ternary complex containing both spacer and complementary protospacer (Abudayyeh et al., 2016; East-Seletsky et al., 2016; Konermann et al., 2018; Smargon et al., 2017; Yan et al., 2018). To understand the molecular basis for nuclease activation, we sought to determine the ternary structure composed of Cas13d bound to both crRNA and its complementary target RNA. To stabilize the ternary complex in a pre-cleavage state, we mutated all four predicted catalytic HEPN domain residues to alanine (R295A/H300A/R849A/H854A). We previously reported that this “catalytically dead” Cas13d (dCas13d) retains the ability to bind both crRNA and target RNA, but cannot cleave ssRNA (Konermann et al., 2018).

We first assembled the dCas13d binary complex, followed by incubation with a 30 nt long target RNA (nt 1-30) that includes a 22-nt (nt 5-26) complementary protospacer, flanked by 4 nt (nt 1-4, 27-30) overhangs at each end (**Figure 1E**). An additional gel filtration step led to a construct that was monodisperse and could be isolated for cryo-EM studies (**Figure S2A-B**). Using similar procedures, we refined 51,885 particles to determine the structure of Cas13d ternary complex to an average resolution of ~3.3 Å, with a range in local resolution distribution that is similar to Cas13d binary complex (**Figure S2C-D**), albeit a more anisotropic distribution of directional resolution (**Figure S2E-G**). The quality and overall features of the map (**Figure 1F**) were sufficient to derive an atomic model of the ternary complex and resolve most of the polypeptide chain, the entire 52 nt crRNA, and all complementary nucleotides of the target RNA (nt 5-26) (**Figure 1G)**. The model is consistent with the cryo-EM density (**Table S1**).

### Cas13d binds target RNA within a large central cleft opposite to the catalytic site

The structure of the Cas13d-crRNA-target-RNA ternary complex shows a similar compact architecture as the binary complex, with the protein subunits of both lobes wrapped around a spacer:protospacer duplex (**Figure 1G**). All 22 complementary nucleotides of the target RNA (nt 5-26) base-pair with the spacer within crRNA **(Figure 1E**), and only the terminal two bps extend outside of the central cleft (**Figure 1F, 1G**). The 5′ handle maintains a solvent-exposed organization as in the binary state, while the guide-target duplex assembles into an A-form RNA helix within the cleft bound by HEPN1, Helical-1, and Helical-2 domains.

On the outside face of the protein, opposite of the central cleft, HEPN1 and HEPN2 form an endoRNase heterodimer. HEPN1 is subdivided in sequence space (residues ~150-344 termed HEPN1-I and ~495-577 termed HEPN1-II), but forms a contiguous tertiary fold. The α1 of HEPN1-I and the C-terminal portion of HEPN2 form the structural backbone of the bipartite active site and position the four catalytic residues (R295A/H300A/R849A/H854A) of the R-X_4_-H motif outward on the external face of Cas13d. This orientation of the HEPN active site primes the Cas13d ternary complex for cleavage of both target and collateral RNAs.

### Bilobed organization is conserved across Class 2 CRISPR effectors

Class 2 CRISPR-Cas effectors are characterized by their bilobed architectures (**Figure S3A-I**) containing a nucleic acid recognition (REC) lobe that binds crRNA, as well as a nuclease (NUC) lobe that is responsible for cleavage of target nucleic acids (Garcia-Doval and Jinek, 2017) (**Figure S3A**). The DR within Cas13a crRNA is buried within protein density, sandwiched between the corresponding NTD and Helical-1 domains (Liu et al., 2017b) (**Figure S3F,G**). In contrast, Cas13d features a compact REC lobe wherein the NTD (residues ~1-150) is responsible for the majority of DR-specific contacts, with the exception of a few residues within HEPN2 (e.g. K932) (**Figure S3H,I**). Notably, the DR within both binary and ternary states prominently protrudes from the effector. A similar compaction, coupled to solvent-exposure of crRNA and spacer:protospacer duplex, has been observed among other small Cas nucleases such as *Sa*Cas9 (Nishimasu et al., 2015) (**Figure S3C**).

### Cas13d maintains an extensive nucleoprotein interface throughout the length of bound RNA

In both its binary and ternary forms, Cas13d makes extensive nucleoprotein interactions with spacer and complementary target protospacer, with all five protein domains contributing to RNA stabilization. Both complexes maintain a similar configuration of the DR, with only minor structural differences between the two forms (**Figure 1D, G**). In the binary complex, most of the length of the 22-nt single-stranded spacer interfaces with key residues of Helical-1, Helical-2, HEPN1, and HEPN2 (**Figure 2)**. The majority of interactions are comprised of backbone contacts, typically with phosphate groups but also ribose hydroxyl groups. Individual base interactions occur at lower frequency and are typically constrained to conserved bases within the DR (**Figure 2 and Figure 3A**). These extensive contacts stabilize the spacer region in a primed pseudo-helical conformation within the solvent-exposed central channel (**Figure 3B-D**).

**Figure 2.**
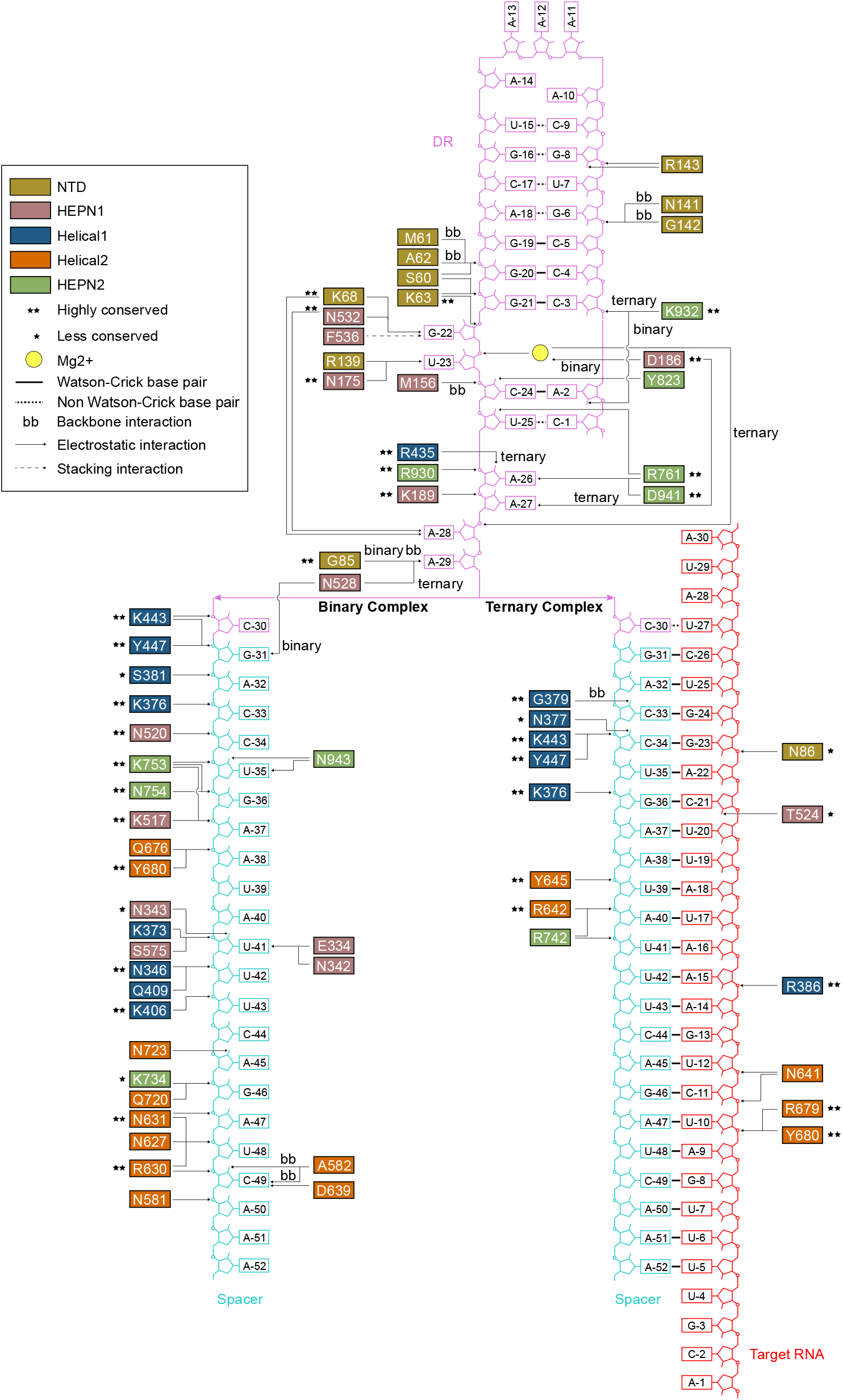
Cas13d nucleoprotein interactions reconfigure between its binary and ternary forms. Schematic of protein:DNA interactions within binary and ternary forms of Cas13d. Arrows point to the specific interactions. Standard amino acid and nucleotide abbreviations apply to all residues and bases. Colors as in Figures 1-2. Conservation of interacting residues and types of interactions are indicated in the legend.

**Figure 3.**
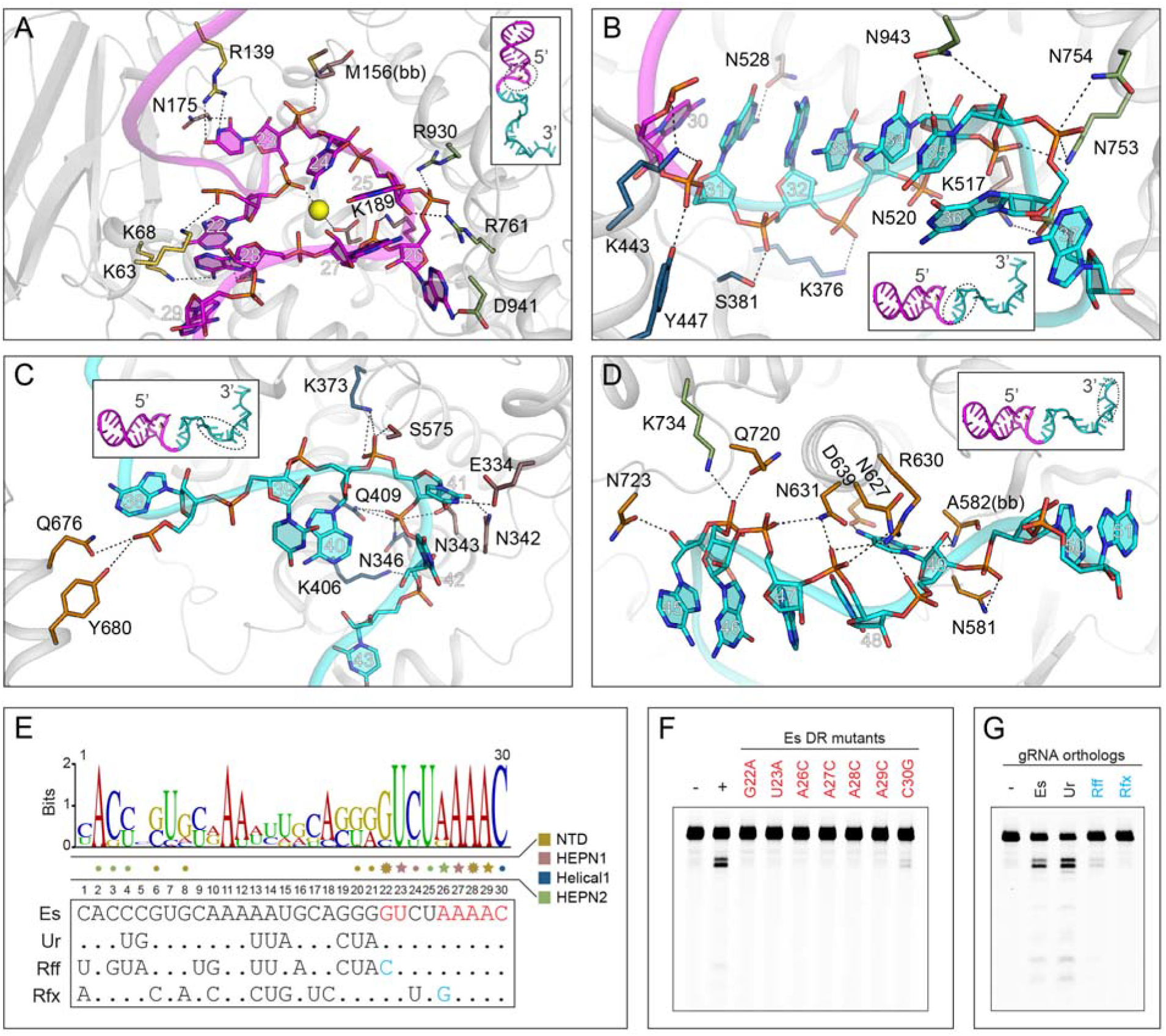
Recognition of crRNA. (A) Close-up view of the nucleoprotein interactions at the DR-spacer interface in the binary complex. (B-D) Close-up view of the nucleoprotein interactions at the (B) 5′ end, (C) middle region, and (D) 3′ end of spacer RNA in the binary complex. (E) processed 30nt DR sequence identity logo among seven previously reported Cas13d orthologs (top) with sequences of four selected orthologs shown (bottom). A black dot indicates an identical residue to *Es*Cas13d. A star above a residue indicates protein:base interactions, and a dot indicates protein:backbone interactions. Mutations used in F) are highlighted in red and critical residue differences between orthologs (for G) are indicated in blue. (F) *Es*Cas13d requires the formation of key protein-base interactions to maintain efficient cleavage of complementary ssRNA targets. Substitutions within the DR that make protein:base interactions (see panels B-D) abolish or compromise cleavage. C30 resides in a complex environment and can potentially make both base- and backbone-specific interactions, thus accounting for only partial loss of activity. (G) Neither the Rff nor Rfx DR can be substituted for the proper Es DR to efficiently cleave complementary ssRNA targets as predicted. Ur DR can trans-complement Es, because the effector:base interactions are maintained (see panel E).

Upon target binding, most of the spacer interactions are rearranged following formation of the A-form dsRNA helix. Only four residues (K443, Y447, K376, Y680) interface with RNA in both enzymatic forms, forming backbone contacts with the 5′ end of the spacer in binary form. Upon target binding, all of these interactions are shifted towards the 3′ end, reflecting spacer compaction protein rearrangements. Overall, interactions along the RNA duplex are sparser in ternary and exclusively confined to the RNA backbone (**Figure 2**).

### Cas13d recognizes the Mg^2+^-stabilized 5′ handle of crRNA

Specific recognition and binding of the constant 5′ handle within their cognate crRNAs is a key requirement for all class 2 CRISPR effectors. In the Cas13d binary structure, we identified multiple residues that interact in either a base- or backbone-specific manner with the DR of the crRNA. (**Figure 2 and Figure 3A-D).** The base-specific contacts are concentrated within the unpaired, conserved terminal nucleotides of the DR motif (nt 22-30) and include G22, U23 from the 2-nt bulge region, as well as A26, A27, A28, and A29 within the 5 terminal nt of the DR (nt 26-30). The 2-nt bulge (nt 22-23) appears to be an invariant feature among Type VI RNA-guided RNases (Abudayyeh et al., 2016; East-Seletsky et al., 2016; Konermann et al., 2018; Shmakov et al., 2015; Smargon et al., 2017; Yan et al., 2018). The 5-nt 5′-AAAAC-3′ motif at the 3′ end of the DR (nt 26-30) is generally conserved across Cas13d orthologs (Konermann et al., 2018), likely owing to its recognition by the Cas1/Cas2 adaptation module for spacer integration (Wright and Doudna, 2016) (**Figure 2 and Figure 3A**). Mutagenesis of the corresponding crRNA nucleotides abolished Cas13d-mediated ssRNA cleavage, confirming the importance of these interactions for proper crRNA binding and positioning (**Figure 3E and F**).

Given the absence of base-specific contacts along the majority of the 5′ handle (nt 1-21), we reasoned that *Es*Cas13d would be able to utilize distinct crRNA of other Cas13d orthologs where the terminal 8 nt of crRNA is conserved. As predicted, *Es*Cas13d maintained full target cleavage activity with the *Ur*Cas13d cognate crRNA, which contains numerous DR mutations relative to the *Es*Cas13d crRNA but maintains the necessary base-specific contacts. In contrast, crRNAs from *Rff*Cas13d and *Rfx*Cas13d were predicted to disrupt the critical base-specific G22 and G26 contacts; accordingly, target cleavage activity was abolished (**Figure 3G**). These data provide a structural basis for defining key base requirements and likely crRNA exchangeability across the Cas13d family, and further studies should elaborate functionally orthogonal subfamilies.

A prominent feature characterizing the nucleoprotein interface in both the binary and ternary forms of Cas13d is the highly ordered, albeit irregularly shaped single-stranded RNA at the 3′ end of the DR (**Figure 2, 3A**). This general region (nt 23-29) forms a hairpin loop surrounding a centrally located Mg^2+^ ion in both binary (**Figure 3A**) and ternary complexes (**Figure 4A**). Two negatively charged phosphates from U23 and A28 tightly coordinate Mg^2+^, with the phosphates of U25 and A27 and the side chains of D186 and N183 in the immediate vicinity, all of which can potentially contribute to metal coordination. Mg^2+^ frequently exploits only two immediate coordination positions by nearby phosphate or carboxylic chemical groups, with the others taken up by water molecules to form an octahedral coordination geometry (Harding et al., 2010).

**Figure 4.**
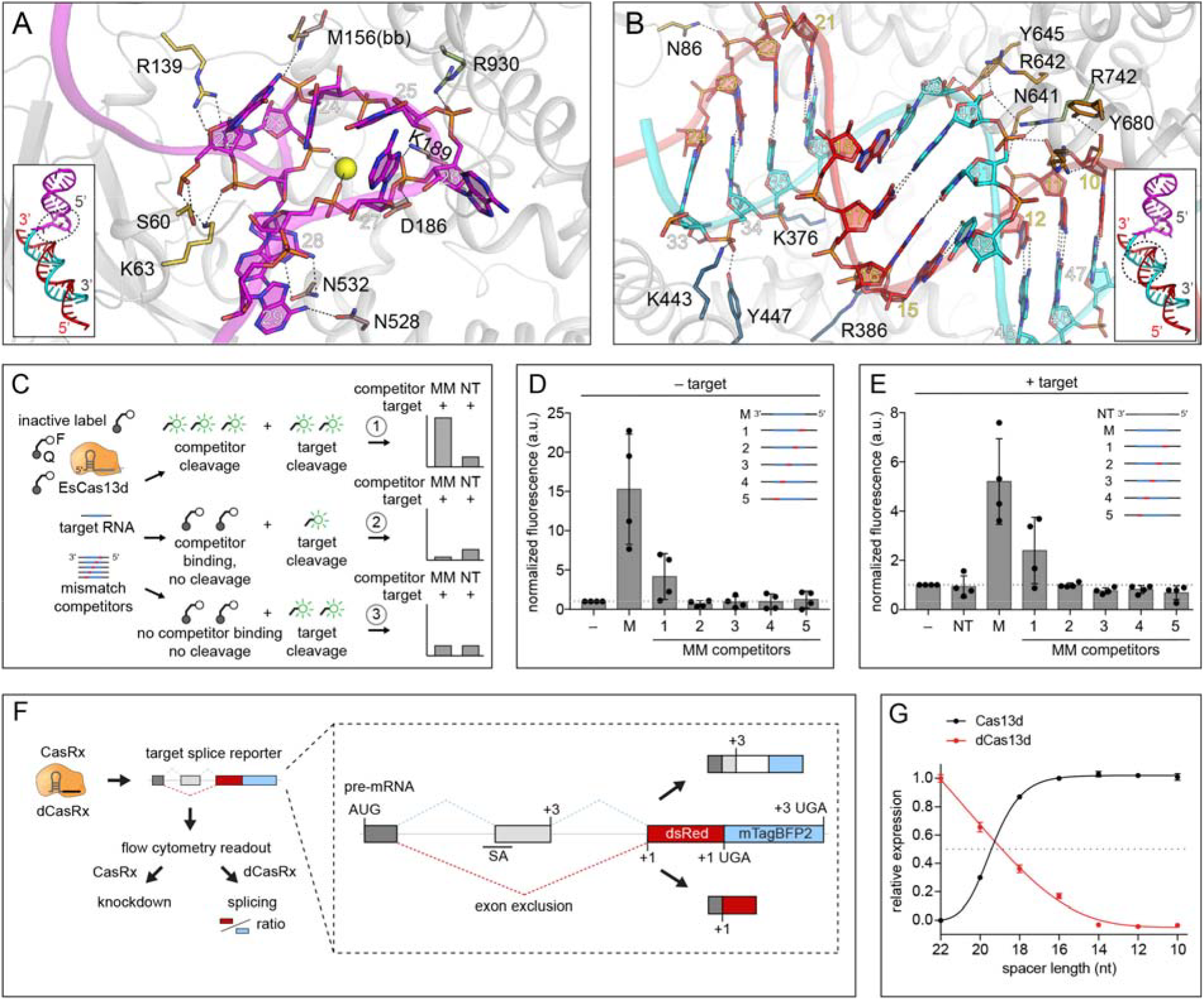
Complementarity requirements for Cas13d ribonuclease activation. (A) Close-up view of the nucleoprotein interactions at the DR-spacer interface in the ternary complex. (B) Close-up view of the nucleoprotein interactions surrounding the spacer:protospacer duplex. (C) Schematic of competitor cleavage assay describing three possible scenarios and expected results. MM, mismatched competitors with 4 nt of non-complementarity to the crRNA. NT, non-targeting competitors with no complementarity to the crRNA. (D) Quantified fluorescent signal of cleavage assay containing Cas13d, crRNA and competitor RNAs only (no matching target). Signal is normalized to a negative control containing Cas13d and crRNA only. M, fully matched competitor. Each condition represents activity across four distinct target sequences averaged from 3 replicates (mean ± SD). (E) Quantified fluorescent signal of cleavage assay containing Cas13d, crRNA, target RNA, and competitor RNA at 25X molar excess to target. Signal is normalized to the condition with Cas13d, crRNA and target without competitor. Each condition represents activity across four distinct target sequences averaged from 3 replicates (mean ± SD). (F) Illustration of parallel readout for Cas13d cleavage and binding in human cells using a fluorescent splicing reporter with the *Rfx*Cas13d-NLS (CasRx) ortholog. (G) CasRx cleavage and dCasRx splicing (binding) activity as a function of crRNA spacer length. Cleavage and splicing were each normalized to a non-targeting crRNA and represented relative to the activity of the full-length (22nt) crRNA (mean ± SD, n = 3).

Although we previously showed that Mg^2+^ is not required for pre-crRNA cleavage by *Es*Cas13d (Konermann et al., 2018), generation of mature crRNA can be inhibited by the addition of EDTA when pre-crRNA is in excess to Cas13d protein (Yan et al., 2018). We found that Mg^2+^ ions increase array processing efficiency at lower Cas13d protein concentrations (**Figure S4**). These data indicate that Mg^2+^ stimulates pre-crRNA cleavage, likely by facilitating proper pre-crRNA folding for efficient Cas13d recognition and processing and/or by increasing Cas13d crRNA binding affinity directly. Mg^2+^ has been shown to have a similar role in increasing Cas12a/Cpf1 affinity for crRNA and stimulating crRNA processing (Swarts et al., 2017), highlighting its role across diverse Class 2 effectors.

While most HEPN domains have a metal-independent endoRNase activity (Anantharaman et al., 2013), Mg^2+^ appears to be essential for Cas13d ssRNA cleavage (Konermann et al., 2018; Yan et al., 2018), and this requirement is conserved across the Type VI family of Cas13 enzymes (Abudayyeh et al., 2016; East-Seletsky et al., 2016; Smargon et al., 2017). Although we cannot exclude the possibility that other Mg^2+^ ions are required to coordinate target cleavage, the crRNA-coordinated Mg^2+^ likely contributes indirectly to target cleavage by stabilizing crRNA interaction with Cas13d, a phenomenon that is found within numerous ribozymes and RNA structures.

### Target duplex formation and lack of PFS requirement

In the active, binary Cas13d surveillance complex, most of the single-stranded spacer is solvent exposed and structurally poised for initiating base-pairing with potential target sequences to form an A-form dsRNA helix in the ternary complex (**Figure 4B**). For DNA-targeting class 2 CRISPR effectors, target interaction is initiated by protein-PAM (protospacer adjacent motif) interactions. Cas13d lacks the analogous protospacer flanking sequence (PFS) requirements (Konermann et al., 2018; Yan et al., 2018). In contrast, some Cas13a orthologs (including *Lsh*Cas13a) have been reported to display a single base 3′ H (non-G) PFS. This was previously proposed to be caused by base-pairing of the terminal conserved C(30) of crRNA DR with a complementary target G, which would then destabilize critical contacts with the HEPN-1 domain following an outward rotation of the C(30) base (**Figure S5A**) (Liu et al., 2017a). In *Es*Cas13d, the C(30) base is already flipped out in its ternary form despite a mismatch with the target base (U) (**Figure S5B**). A complementary G nucleotide would therefore not be expected to cause any additional rearrangement, suggesting a structural rationale for the absence of a PFS requirement in Cas13d.

### Cas13d binding and cleavage are interlinked

Given the lack of an overt PFS requirement, we sought to understand Cas13d target binding and cleavage in the context of target complementarity. Class 2 CRISPR-Cas nucleases exhibit distinct mechanisms for target binding and cleavage. Cas9 from *Streptococcus pyogenes* exploits a sequential target binding mechanism characterized by stepwise PAM-proximal strand invasion. Stable binding is observed after ~12 nt of complementarity based on both *in vitro* (Sternberg et al., 2014) and cell data (Dahlman et al., 2015; Kiani et al., 2015), while mismatches in this “seed” region are poorly tolerated (Hsu et al., 2013). In contrast to binding, Cas9 target cleavage requires ~4 nt additional matches to facilitate proper rotation of the HNH catalytic domain (Sternberg et al., 2015). In contrast to *Sp*Cas9, Cas12a/Cpf1 family proteins require extended complementarity of at least 17 nt for stable binding (Singh et al., 2017). Cas13 binding and cleavage requirements are largely unclear, although a central seed region has been proposed for both Cas13a and Cas13b based on the observation that mismatches are least tolerated in the center of the spacer (Abudayyeh et al., 2016; East-Seletsky et al., 2016; Knott et al., 2017; Liu et al., 2017b; Smargon et al., 2017).

To investigate Cas13d target binding and cleavage complementarity, we conducted cleavage assays with a panel of target RNA competitors carrying 4-nt mismatches at different positions along the target sequence (**Figure 4C**). To control for possible spacer sequence-intrinsic effects, we generated competitors for four distinct crRNA:target pairs. In this assay, incubation of Cas13d-crRNA with a complementary target drives trans-ssRNA cleavage of a quenched fluorescent reporter bystander, releasing a detectable fluorescent signal. We sought to distinguish between three possible scenarios when a competitor RNA is introduced at high molar excess relative to the target RNA: Cas13d 1) directly cleaves competitor, increasing fluorescent signal relative to target alone, 2) binds competitor without forming a catalytically active ternary complex, thereby sequestering Cas13d from target RNA and decreasing fluorescent signal, or 3) is unable to bind or cut competitor, with no resulting change in fluorescence.

In the absence of target, perfectly matched competitors triggered a robust increase in fluorescence as expected (**Figure 4D**). Competitors carrying 5′-proximal mismatches activated bystander cleavage in 2/4 cases, indicating that complementarity in the DR-distal region of the crRNA spacer is not strictly required for ternary activation (scenario 1). Bystander activity was not activated by other competitors (scenario 3), suggesting a lack of a consistent seed region within the crRNA spacer for target cleavage. In the presence of target, we observed a similar pattern of Cas13d activity across all competitor mismatch positions to the target-free condition (**Figure 4E)**. In particular, none of the mismatch competitors mediated a decrease in fluorescence (in contrast to scenario 2), suggesting that stable target binding requires at least 18 nt of complementarity and that binding and cleavage are closely coupled in Cas13d.

To further explore the interdependence between nucleotide complementarity and cleavage efficiency, we tested a closely related Cas13d ortholog from *Ruminococcus flavefaciens* XPD3002 (*Rfx*Cas13d) in a cell-based reporter to assay for Cas13 knockdown and splicing as a proxy for binding and cleavage activity (Konermann et al., 2018) (**Figure 4F**). Using a series of crRNA spacer truncations progressing from 22-nt to 10-nt, we observed a simultaneous decrease in knockdown and splicing as a function of decreasing spacer length (**Figure 4G**). Half-maximal activity occurred between 18 and 20-nt spacer length.

Taken together, the lack of a clear seed is consistent with the observation of a solvent exposed spacer stabilized in an accessible configuration along the open channel in the binary complex, positioning crRNA to initiate base-pairing with protospacer at multiple possible locations. Beginning with ~18 nt of complementarity, Cas13d likely undergoes a partial reconfiguration of its relevant nucleoprotein interface. Upon reaching complete 22 nt guide-target duplex complementarity, the enzyme transitions to a maximally active and cleavage-competent state.

### Target RNA binding reconfigures Cas13d into an ssRNA cleavage complex and allosterically activates the HEPN domains

The transition from the binary surveillance complex to the ternary complex activates Cas13d for RNA cleavage. Numerous conformational rearrangements occur during this transition, stabilizing the activated cleavage-competent state within the catalytic HEPN domain dimer. HEPN domains function as obligate dimers, with 2 R-X_4-6_-H motifs forming a bipartite active site to mediate RNA hydrolysis (Anantharaman et al., 2013). While dimerization is required for HEPN activity, the final active conformation is often triggered by distinct ligands, such as cyclic oligoadenylate for Csm6 (Kazlauskiene et al., 2017; Niewoehner et al., 2017), 2-5A and ADP for RNase L (Huang et al., 2014), or crRNA-mediated binding of target RNA by Cas13 (Liu et al., 2017a).

Cas13d target binding triggers the most dramatic polypeptide rearrangements within the Helical-1 domain, which shifts outward by an average Cα-Cα RMSD of ~12 Å to accommodate complete target RNA binding within an expanded cleft. To a lesser extent, portions of the HEPN1, Helical-2, and HEPN2 domains, particularly regions proximal to the 3′ end of spacer (5′ end of protospacer), all undergo subtler structural changes (**Figure 5A-B**). Whereas most of the DR remains unperturbed, the majority of the spacer reorganizes from a single-stranded pseudo-helical conformation into a double-stranded A-form RNA helix (**Figure 5C-D**). Collectively, such rearrangements widen the RNA-binding cleft, ranging from ~13Å at the narrowest point between NTD and Helical-1 (**Figure 5E**) and from 12Å to 44Å between Helical-1 and HEPN2 at the 3′ end of crRNA (**Figure 5F, Movie S1)**.

**Figure 5.**
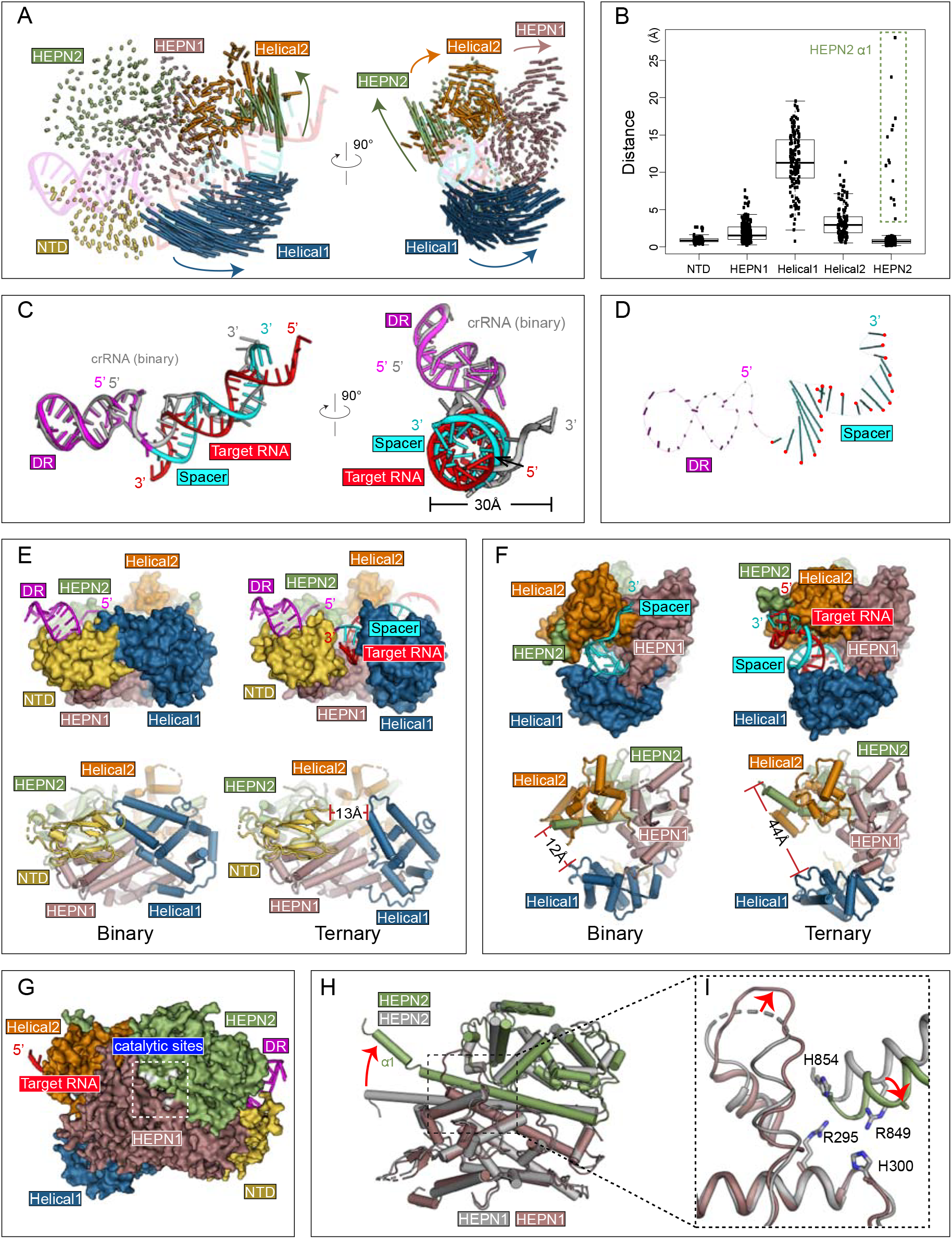
Conformational rearrangements of Cas13d upon target substrate recognition. (A) Schematic representation of Cas13d domain movement. Each vector represents a Cα atom translation from its binary to ternary forms. Vector length corresponds to the translational distance. The directions of movement are indicated by arrows. (B) Quantitative measurement of Cas13d domain movement. The residues that undergo largest translations in HEPN2 are highlighted with a dashed box. (C) Superposition of the crRNA in the ternary complex (DR region in magenta and spacer in cyan) with that in the binary complex (in gray). (D) Schematic representation of Cas13d crRNA movement. Vector representations as in (A). The position of the phosphate atom in the ternary structure are highlighted in red. (E-F) Target RNA binding widens the central channel and (E) opens the cleft between NTD and Helical-1 and (F) increases distance between Helical-1 and HEPN2. (G-I) Stabilization of the HEPN catalytic domain dimer. (G) Surface representation of Cas13d ternary complex, with HEPN catalytic residues highlighted in white. (H) Superposition of HEPN1 and HEPN2, and (I) a close-up view of the catalytic center. The four catalytic residues advance inward and a nearby loop is stabilized upon target binding.

We also observe some intra-domain reorganization within Cas13d (**Figure S6A**). Most prominently, a kink is introduced into the C-terminal region of HEPN1-1, while the N-terminal loop of HEPN1-II (and, correspondingly, the C-terminal loop residues of Helical-1) reposition by an average Cα-Cα RMSD of ~4 Å. Both of these correspond to the two flexible linkers connecting Helical-1 and, together, these rearrangements are responsible for repositioning Helical-1 as it cradles incoming target RNA (**Figure S6B-C**). Further, α1 of HEPN2 bends into two pieces (α1-1, α1-2) in addition to numerous local rearrangements within Helical-2 that collectively amount to an average Cα-Cα RMSD of ~4.5 Å across this domain (**Figure S6A**).

The HEPN2 domain, which resides on the “back side” of Cas13d (**Figure 5G**), undergoes several rearrangements to facilitate target cleavage. A structural alignment of HEPN1-I between binary and ternary forms indicates the catalytic residues of HEPN2 (R849 and H854) reposition by ~4 Å, shifting closer to the corresponding catalytic residues of HEPN1-I (R295 and H300) (**Figure 5H-I**). In turn, loop 204-214 of HEPN1-I, which is poorly ordered and unmodeled in the binary form, leans up against HEPN2, forming numerous salt bridges that presumably stabilize the re-organization and activation of the catalytic site. In sum, given that target RNA resides on the opposite side of the HEPN catalytic site, these rearrangements suggest allosteric HEPN activation modulated by target RNA binding. Together, the structural changes serve to (1) accommodate target RNA binding and (2) reconfigure the catalytic site into its cleavage-competent form.

### Apo Cas13d utilizes multiple dynamic domains to form an RNA-binding cleft that is stabilized upon guide binding

To better understand the mechanism of crRNA binding, we examined the apo form of Cas13d. We used similar strategies for purifying the sample, preparing cryo-EM grids, data collection, and image analysis. A large data set provided 330,986 particles, of which 154,889 were selected for 3D classification and refinement (**Figure S7A**). We observed variable density within the 2D class averages (**Figure S7B)**, in contrast with the sharp signal present for both the binary and ternary data sets (**Figure S1B, S2B**). Despite the apparent heterogeneity, an *ab initio* 3D reconstruction led to a map that was refined to ~6.5 Å resolution (**Figure 6A-B, Figure S7C-G, and Supplementary Table 1**). Strikingly, the apo Cas13d reconstruction accounts for only part of the mass in comparison to both binary and ternary forms, with the majority of homogeneous density corresponding to a stable α-helical core (**Figure 6C**). To verify that the remaining density was present in the data, we derived 2D class averages *ab initio* and found that many of these captured protein density that was otherwise unresolved in the 3D reconstruction (**Figure S7H**).

**Figure 6.**
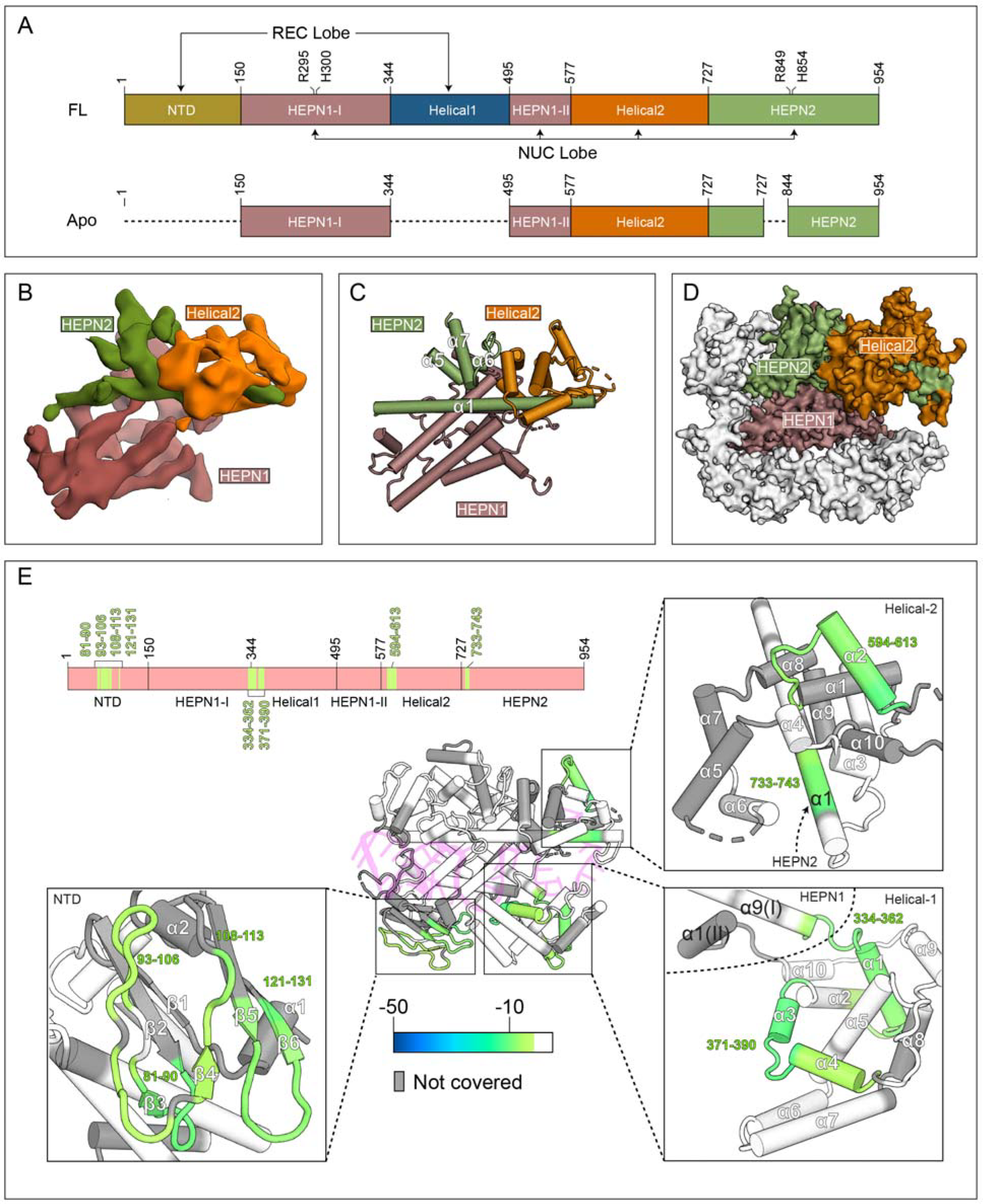
Apo Cas13d contains mobile domains that are stabilized upon RNA binding. (A) Domain organization of full-length *Es*Cas13d and the visible portion of apo *Es*Cas13d by cryo-EM (see below). Domain boundaries and catalytic sites of the HEPN domains (R295, H300, R849, H854) are indicated. Undetectable regions in the apo Cas13d reconstruction (see B-D) are represented with dashed lines (B) Cryo-EM reconstruction of apo *Es*Cas13d, resolved to 6.5 Å resolution. (C) Rigid-body docked model of *Es*Cas13d based on the cryo-EM density. (D) Superposition of the structured portion of apo *Es*Cas13d evident within the cryo-EM reconstruction (surface representation) with that of *Es*Cas13d in the binary form. (E) Differential HDX colored onto the binary protein form of Cas13d. Regions that exhibit statistically significant changes in amide exchange are colored onto the structure, and onto a domain organization (top). Insets show four distinct close-up views. White indicates no statistically significant difference; gray indicates no coverage in the differential HDX map.

We could readily assign the known domain boundaries by docking the binary model into the apo reconstruction. Portions of HEPN1, Helical-2, and HEPN2 (NUC lobe) account for the mass derived by cryo-EM, whereas the entire REC lobe (NTD and Helical-1), as well as small surrounding regions of the NUC lobe, were invisible (**Figure 6D**). The domain organization of NUC lobe in the apo enzyme remains largely unchanged relative to the binary complex, at least at the nominal ~6.5 Å resolution. These data suggest that the REC lobe (NTD, Helical-1) and portions of HEPN2 may be dynamically arranged in the absence of RNA. Further, because cryo-EM is performed under solution conditions, the observed dynamics is likely an inherent property of the enzyme.

To understand protein conformational dynamics at the sequence level, we conducted hydrogen-deuterium exchange (HDX) and compared the stability of individual regions of Cas13d between its apo and binary states. HDX measures the rate of deuterium incorporation into backbone amides under solution conditions and, when paired with protease digestion and mass spectrometry, provides a direct readout of deuterium incorporation into individual peptides (Englander, 2006). Because unstructured and flexible regions of Cas13d undergo deuterium exchange more rapidly than those that are stable and hydrogen-bonded, HDX can directly probe for conformational dynamics on a residue-by-residue level. We subjected both the apo and the binary forms to HDX-MS to generate both their individual exchange profiles as well as a differential exchange map (**Figure S8**).

Within independent experiments containing either apo or binary forms of Cas13d, individual exchange profiles showed similar patterns at the sequence level (**Figure S8A, B**), indicating that the constituent domains, including the REC lobe, exhibit similar overall folds. However, several regions of Cas13d were clearly stabilized upon RNA binding, as indicated by reduced exchange in the differential profiles (**Figure 6E**). Three of these regions bind distinct segments within crRNA. Residues ~81-131 form part of the NTD, where residues K68, G85 and R139 make critical base-specific interactions with the 3′ end of the DR (**Figure 2, Figure 3F-G**), while residues ~371-390 and ~594-613 form an RNA-binding interface with the phosphate backbones of the 5′ and 3′ ends of spacer RNA, respectively. Residues ~733-743 of HEPN2 α1 are missing from the apo EM density and exhibit reduced solvent exchange, suggesting they are also stabilized in the binary complex, with K734 contacting the phosphate backbone between nt 45-46 of the crRNA spacer. Finally, residues ~325-360 represent the interface between HEPN1-I and Helical-1, with residues 335-340 forming a hinge-like structure connecting Helical-1 to the NUC lobe (**Figure S6B-C**).

Taken together, these data indicate that both Cas13d lobes are appropriately folded in the apo configuration and the REC lobe is likely to be mobile relative to the NUC lobe. This is supported by the 2D class averages, which indicate the presence of a second lobe, as well as HDX analysis indicating increased mobility of the NTD domain and the linker region (residues ~325-360) connecting Helical-1 to the NUC lobe. Upon RNA binding, the Cas13d binary complex is stabilized in multiple regions, and a central, positively charged RNA binding cleft is formed between the REC and NUC lobes (**Figure S8C**). Analogous reconfigurations of NUC and REC lobes upon crRNA binding and formation the central positively charged cleft in the binary complex have been reported for other Class 2 CRISPR enzymes, including Cas9 (Jinek et al., 2014; Nishimasu et al., 2015). Furthermore, there is precedent for the observed conformational dynamics of Cas nucleases in their apo forms, especially for Cas12a/Cpf1 (Dong et al., 2016) and possibly for the NTD of Cas13a (Liu et al., 2017b). Our experiments under solution conditions suggest that the REC lobe adapts multiple conformations relative to the NUC lobe in the Cas13d apo state, which may have implications for sampling distinct RNA features and facilitating efficient crRNA recognition and binary transition.

### Cas13d truncations for flexible AAV packaging

The compact size of Cas13d is accompanied by an integration of multiple distinct functions into each individual constituent domain. Recognition of the DR, for example, is achieved by the Helical-1 and NTD domains in *Lbu*Cas13a through extensive protein:RNA interactions (Knott et al., 2017; Liu et al., 2017b). In contrast, Cas13d completely lacks an equivalent to the Helical-1 domain of Cas13a and its structural role is substituted by the helices in HEPN2 that are proximal to the DR (**Figure 2**). In addition to its role in DR recognition (**Figure 3A**) and target cleavage (**Figure 5G-I**), HEPN2 forms stabilizing interactions with the single-stranded spacer (**Figure 2** and **Figure 3B, 3D**). Each of the other four protein domains within Cas13d similarly contribute key protein:RNA contacts in addition to their structural and catalytic functions (**Figure 2**), and a majority of these residues are conserved among Cas13d orthologs (**Figure S9**). As a consequence, short regions of high conservation between Cas13d orthologs are dispersed throughout the linear protein sequence and separated by only short stretches of low conservation **(Figure S9**).

We predicted that all five domains of Cas13d would be essential for its RNase activity, in contrast to our previous demonstration that the REC2 domain of *Sp*Cas9 is largely dispensable for target DNA cleavage (Nishimasu et al., 2014). To evaluate this hypothesis, we designed 6 deletions in the closely related *Rfx*Cas13d ortholog (CasRx) distributed throughout the HEPN1, Helical-1, Helical-2 and HEPN2 domains and evaluated the ability of these truncation mutants to knockdown a fluorescent reporter in human cells (**Figure 7A and B**). In addition to a full Helical-1 deletion (Δ4), five other deletions were focused on smaller external regions of the protein that exhibited low overall conservation and did not contain any identified conserved RNA:protein contacts (**Figure 7B** and **S9**). Despite these selection criteria, only one deletion in Helical-2 (Δ3) exhibited full activity relative to the wild-type enzyme. Δ3 is located entirely on the external surface of Cas13d and avoids the removal of highly conserved residues, while the other deletions contained at least one highly conserved residue and were not tolerated. To assess if any of the five deletions abolishing Cas13d cleavage activity could maintain target binding, as demonstrated in Cas9 (Sternberg et al., 2015), we tested their capability to bind the pre-mRNA of a bichromatic splicing reporter to mediate exon exclusion (**Figure 4F**). All five deletions, including truncations of the least conserved regions of HEPN1-I and HEPN2, exhibited strongly reduced splicing activity compared to full-length dCas13d (**Figure S10A**). Consistent with spacer truncation mutants that simultaneously impair Cas13d binding and cleavage, (**Figure 4C-D**) these data suggest that target binding and cleavage may be closely coupled in Cas13d.

**Figure 7.**
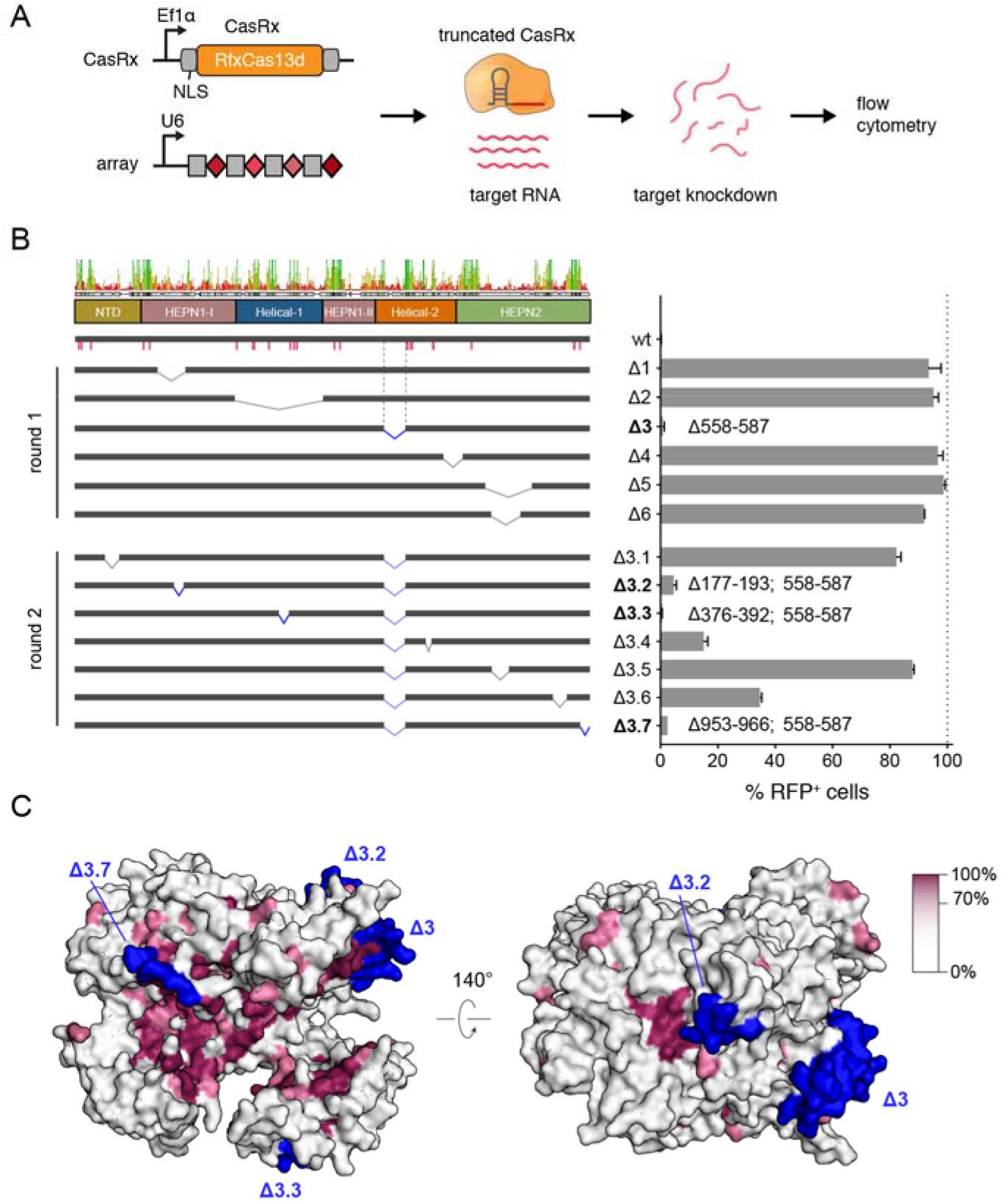
Cas13d truncations and protein conservation. (A) Schematic of the reporter assay used to test CasRx deletion variants in human cells. (B) Knockdown activity of CasRx deletions. Left panel, consensus sequence of the multiple sequence alignment of Cas13d orthologs (**Figure S8**) with domain boundaries and sequence identity as indicated. Pink bars denote conserved protein:crRNA interactions. Deletion junctions are denoted as a linked interval. Knockdown activity is represented relative to non-targeting guide (mean ± SD, n = 3). Deletions exhibiting > 95% reduction of RFP^+^ cells are bolded and the corresponding deletion junction is marked in blue. (C) Surface view of the *Es*Cas13d ternary complex colored by conservation percentage. All truncated regions retaining high knockdown activity (blue) are located on external surface of Cas13d with low (<40%) conservation. Regions with high degree of conservation are concentrated in the RNA-binding cleft (left panel) as well as the HEPN active site (right panel).

Based on our observations from the first round of deletions, we performed a second round of 7 small, surface-localized deletions on top of Δ3 to generate three additional variants with >95% knock-down activity (Δ3.1, Δ3.3 and Δ3.7, **Figure 7B**). The most active resulting variant (Δ3.3) preserves wild-type levels of activity and removes 50AA facilitating AAV-mediated CasRx delivery with increased flexibility. For example, the additional 150bp gained allow for the incorporation of the commonly used WPRE post-transcriptional element within the 4.7 kb AAV packaging limit to improve transgene expression levels (**Figure S10B**).

Furthermore, the identification of four distinct areas of the Cas13d protein amenable to modifications (**Figure 7C**) will aid future engineering efforts, including split CasRx variants or internal insertions of functional domains within the Cas13d ribonucleoprotein complex.

## Discussion

The structural, biochemical, and functional analysis of Type VI *Es*Cas13d presented here reveals 3 distinct states of the Cas13d enzyme as it transitions from its inactive apo (Cas13d) to a surveillance (Cas13d-crRNA) and cleavage-competent (Cas13d-crRNA-target RNA) form.

Cas13d is among the smallest CRISPR-Cas single effectors, with 20-30% less mass than other Type VI Cas13 endoRNases (Konermann et al., 2018; Yan et al., 2018). Compared to Cas13a (Knott et al., 2017; Liu et al., 2017a; Liu et al., 2017b), Cas13d compacts the analogous NTD and Helical-1 domains of Cas13a into a single, 150 aa NTD (**Figure S3**) within the REC lobe of Cas13d. Our data suggests a model whereby REC lobe dynamics within Cas13d may facilitate scanning for the crRNA, and Mg^2+^ cations assist or stabilize appropriate crRNA folding to facilitate efficient effector recognition. Despite the apparent magnesium-dependence of HEPN-mediated target degradation by Cas13d and other Type VI RNases, the DR-bound cation in our structures resides entirely on the opposite side of the ribonucleoprotein complex from the HEPN catalytic site. Mg^2+^ may thus be required for stable formation of a binary complex as a prerequisite for target recognition.

Upon satisfying multiple base-specific contacts within the 3′ end of the DR, crRNA binding triggers stabilization of the binary complex and formation of a central positively-charged cleft between the REC and NUC lobes of Cas13d. The solvent-exposed, single-stranded spacer region takes on a stabilized pseudo-helical conformation which appears to be poised for target binding at multiple positions. Stable RNA duplex formation requires ~18 nt of complementarity and triggers large conformational rearrangements primarily in Helical-1 and HEPN2 to activate the bipartite HEPN domain. Unlike some other Type VI RNases, Cas13d does not require any PFS for target recognition, possibly mediated by the flipped C(30) base within crRNA that avoids base-specific pairing with target RNA. Upon Cas13d ternary formation, the HEPN catalytic residues within the HEPN1 and HEPN2 domains migrate toward one another to generate an external-facing active site. This facilitates direct RNA hydrolysis of both guide-complementary activator RNA and non-complementary bystander RNA for antiviral defense.

Despite their distinct evolutionary origins, functions, and constituent domains, class 2 CRISPR-Cas nucleases share common traits for nucleic acid sensing to confer a host with adaptive immunity. These enzyme types all require structural pre-organization of a guide RNA (crRNA) within a positively-charged protein cleft, coordinated rearrangement into a cleavage complex upon cognate protospacer recognition, and the target-dependent activation of catalytic sites for nucleic acid cleavage (Garcia-Doval and Jinek, 2017). Further, they generally maintain a bilobed architecture, described as a REC lobe for crRNA recognition and a NUC lobe for cleavage, yet CRISPR-Cas single effectors vary substantially in size, shape, domain architecture, and organization.

Within individual Cas types (e.g. Type VI), there is often minimal sequence conservation across subtypes. Cas13d enzymes, for example, do not share sequence homology with Cas13a apart from the minimal 6 aa HEPN catalytic motifs (Konermann et al., 2018; Yan et al., 2018), despite overall similarity of their RNase activities (Abudayyeh et al., 2016; East-Seletsky et al., 2017; East-Seletsky et al., 2016; Konermann et al., 2018). Independent origins of type V subtypes from mobile genetic elements, as previously suggested for DNA-targeting type II and V effectors and subtypes via distinct TnpB transposase subfamilies (Koonin et al., 2017), may explain the convergence and divergence of Cas13 superfamily function and structural organization.

Overall, our data elucidates the structural basis of Cas13d RNA-guided RNase activity and its compaction of these properties into a minimal effector size relative to other Type VI and Class 2 CRISPR effectors, providing a blueprint for improving Cas13d-based RNA targeting tools. Further engineering of smaller Cas13d variants, as shown here, will enable flexible packaging into size-limited viral vectors with large regulatory elements for optimal transgene expression and activity (**Figure S10**). Furthermore, base-specific contacts of Cas13d with the 5′ handle of crRNA were sufficient to delineate crRNA exchangeability across distinct Cas13d orthologs, defining functionally orthogonal subfamilies that could be exploited to facilitate Cas13-based multiplexing applications in both cellular (Abudayyeh et al., 2017; Konermann et al., 2018) and cell-free systems (Gootenberg et al., 2017). Some Cas13d orthologs have accessory proteins that enhance activity (Yan et al., 2018) and could provide clues for improving Cas13d binding or cleavage. In analogy to engineered variants of Cas9 and related nucleases, structure-guided engineering of diverse Cas13d enzymes can be expected to enable improved properties for diverse biomolecular applications of RNA targeting.

## Acknowledgements

We thank Bill Anderson at TSRI and Youngmin Jeon at Salk for help with EM data collection, and Bruce D. Pascal for assistance in analysis and rendering of HDX data. We also thank Tony Hunter, as well as members of the Hsu and Lyumkis labs for critical reading of the manuscript. This work was supported by the Howard Hughes Medical Institute Hannah H. Gray Fellows program (to S.K.), NIH DP5 OD021369 (to P.D.H.), NIH DP5 OD021396 and U54GM103368 (to D.L.), and the Helmsley Charitable Trust (to P.D.H. and D.L.).

## Accession codes and deposition

The electron density maps for apo, binary, and ternary Cas13d will be deposited into EMDB. The models for the binary and ternary complexes will be deposited into the PDB.

## Author contributions

C.Z. performed all cryo-EM data collections, image analysis, model building and refinement. S.K. and P.D.H. purified samples and S.K. generated the complexes. S.K., N.J.B. P.L., and P.D.H. performed the biochemical analyses and genetic analyses. S.J.N. and T.S. performed HDX analyses. P.R.G. supervised HDX studies. S.K., P.D.H. and D.L. conceived this project and P.D.H. and D.L supervised studies. C.Z., S.K., P.D.H., and D.L. interpreted the results and wrote the manuscript.

---

**Figure S1.**
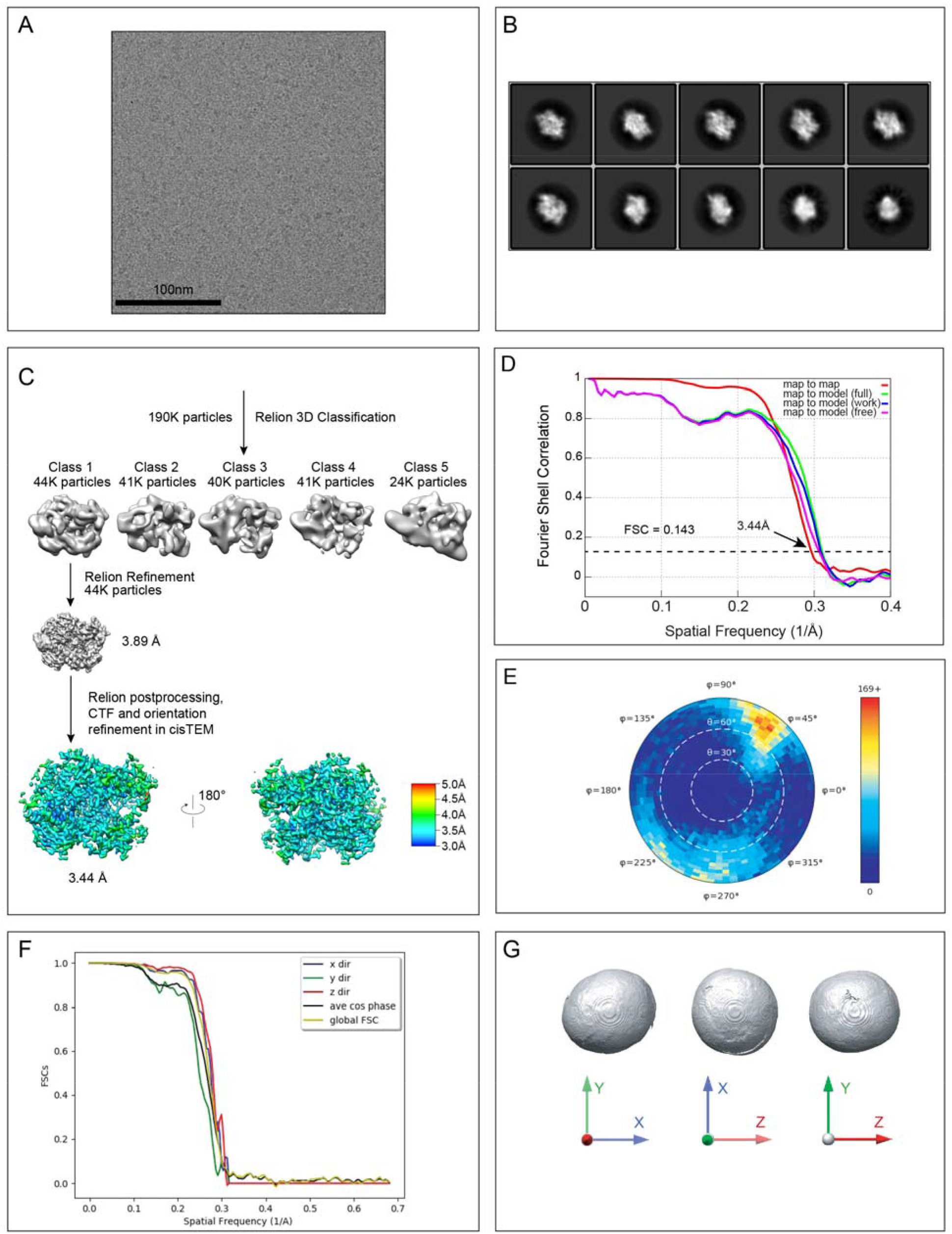
Cryo-EM data collection, analysis, and modeling of Cas13d-crRNA binary complex, Related to Figure 1. (A) Representative cryo-EM micrograph of Cas13d-crRNA binary complex after motion correction. (B) Representative 2D class averages calculated using Relion. (C) Classification strategy and refined maps of Cas13d-crRNA complex, colored by local resolution, calculated using “sxlocres.py” implemented within Sparx. The central RNA-binding groove is characterized by a higher resolution (closer to ~3 Å) than the peripheral regions of the protein (closer to ~4 Å). (D) Fourier Shell Correlation (FSC) curves for cross-validation between the maps and models. Curves calculated between two half maps (red), between the model and the working half map used for model refinement (blue), between the model and the free half map used for validation (magenta), and between the model and the full map (green). (E) Euler angle distribution plot showing the relative orientation of the particles used in the final 3D reconstruction. (F) 3D FSC curves along z (red), y (green) and x (blue) axes, as well as the global FSC (yellow) and the cosine of the phase residual curves (black) are displayed. (G) 3D FSC isosurfaces, thresholded at a cutoff of 0.75 and displayed in three axial orientations, describe the directional resolution (isotropy) of the refined map.

**Figure S2.**
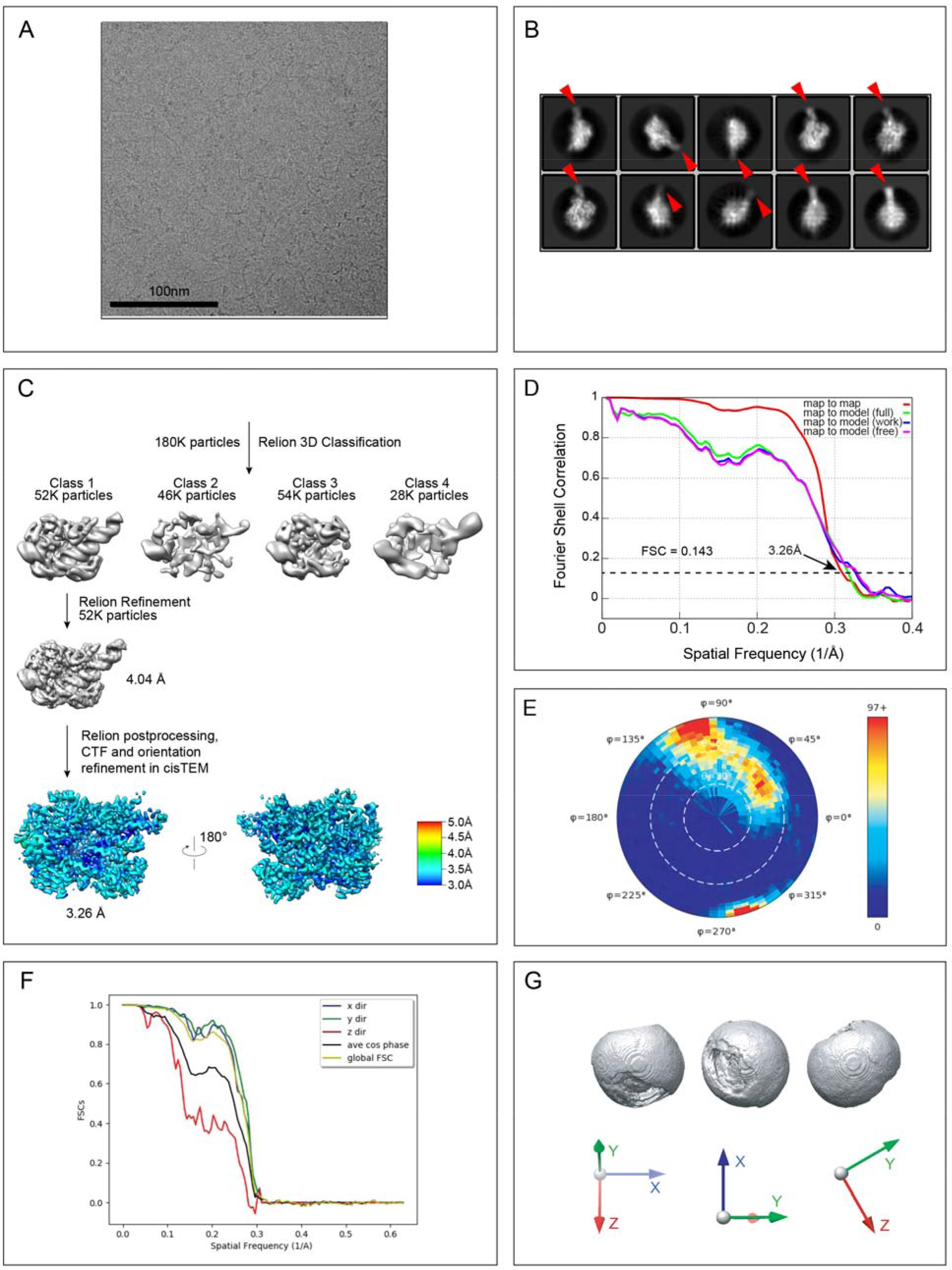
Cryo-EM data collection, analysis, and modeling of Cas13d-crRNA-target-RNA ternary complex, Related to Figure 1. (A) Representative cryo-EM micrograph of Cas13d-crRNA-target-RNA ternary complex after motion correction. (B) Representative 2D class averages calculated using Relion. We observe elongated RNA density downstream of the crRNA (red arrow), which is caused by binding of an additional copy of free target RNA to a palindromic sequence at the 5′ overhang of crRNA, and which becomes progressively disordered further out away from the core density in the reconstructed map. (C) Classification strategy and refined maps of Cas13d-crRNA-target-RNA ternary complex colored by local resolutions, calculated using “sxlocres.py” implemented within Sparx. A higher local resolution is observed near the buried regions, in the vicinity of the RNA, similar to the binary complex (**Figure S1C**). (D) Fourier Shell Correlation (FSC) curves for cross-validation between the maps and models. Curves calculated between two half maps (red), between the model and the working half map used for model refinement (blue), between the model and the free half map used for validation (magenta), and between the model and the full map (green). (E) Euler angle distribution plot showing the relative orientation of the particles used in the final 3D reconstruction. (F) 3D FSC curves along z (red), y (green) and x (blue) axes, as well as the global FSC (yellow) and the cosine of the phase residual curves (black) are displayed. (G) 3D FSC isosurfaces, thresholded at a cutoff of 0.75 and displayed in three axial orientations, describe the directional resolution (isotropy) of the refined map.

**Figure S3.**
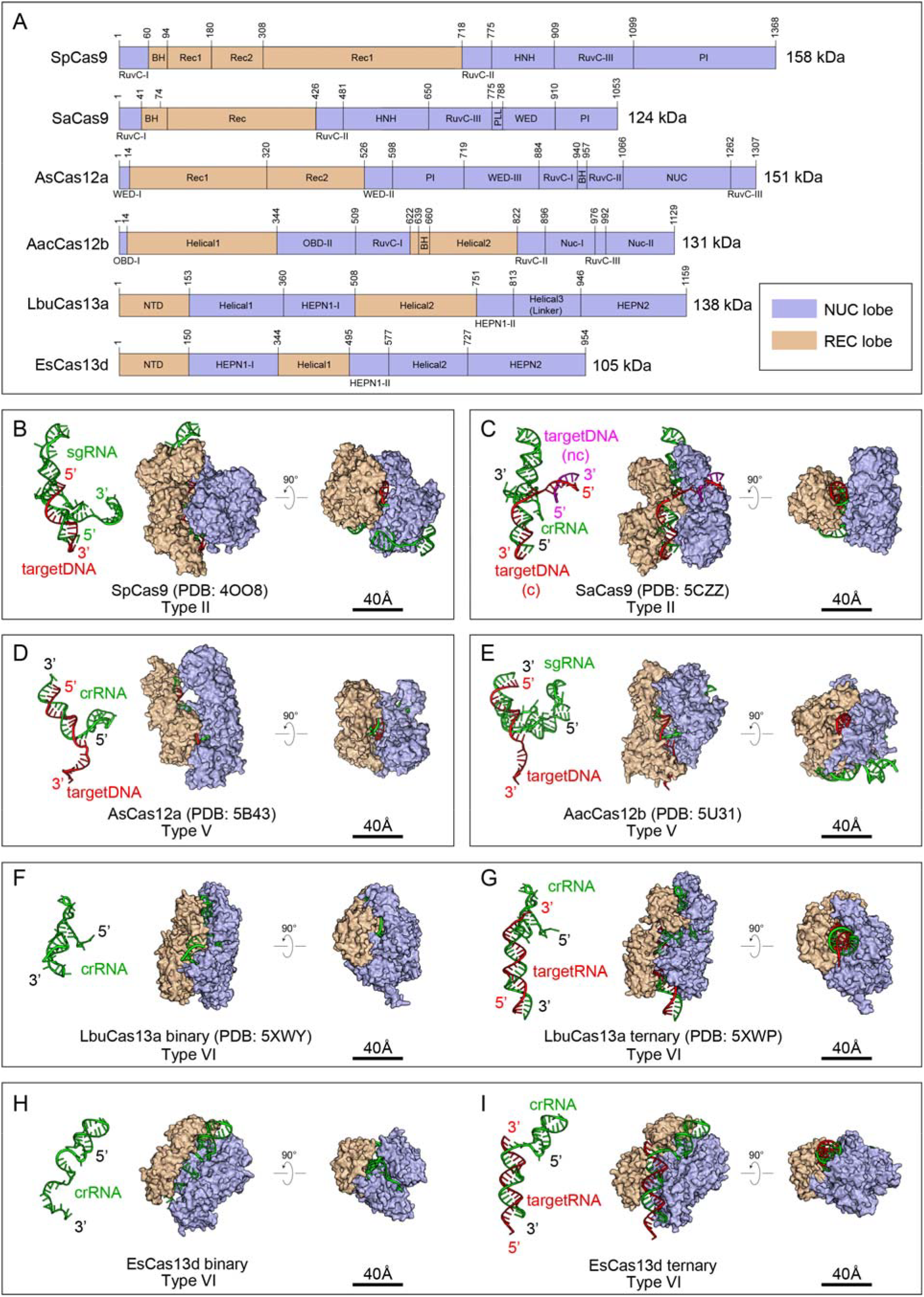
Bilobed structures of representative Class 2 CRISPR effectors. (A) Domain organizations of Class 2 CRISPR effectors: SpCas9, SaCas9, AsCas12a, AacCas12b, LbuCas13a and EsCas13d. The Helical-1 domain of Cas13a has been redefined here to include Cas13 regions The REC and NUC domains are colored wheat and light blue, respectively. (B-I) Surface representations of crystal structures of (B) SpCas9 (PDB: 4OO8), (C) SaCas9 (PDB: 5CZZ), (D) AsCas12a (PDB: 5B43), (E) AacCas12b (PDB: 5U31) (F) cryo-EM structure of LbuCas13a in the binary form (PDB: 5XWY), (G) crystal structure of LbuCas13a in the ternary form (PDB:5XWP), (H) *Es*Cas13d in binary form, (I) *Es*Cas13d in ternary forms. Nucleic acid components and orthogonal views of each structure are shown. The REC lobe of LbuCas13a was proposed to include NTD, Helical-1, and Helical-2, and the NUC lobe to include HEPN1, Helical3/Linker and HEPN2 domains. In our representation, to be consistent with grouping on each side of crRNA, we defined the REC lobe of LbuCas13a as including NTD and Helical-2, whereas the NUC lobe would contain Helical-1, HEPN1, HEPN2, and Helical3/Linker. In *Es*Cas13d, The NTD and Helical1 domains are grouped into the REC lobe, and its NUC lobe contains HEPN1, Helical-2 and HEPN2 domains.

**Figure S4.**
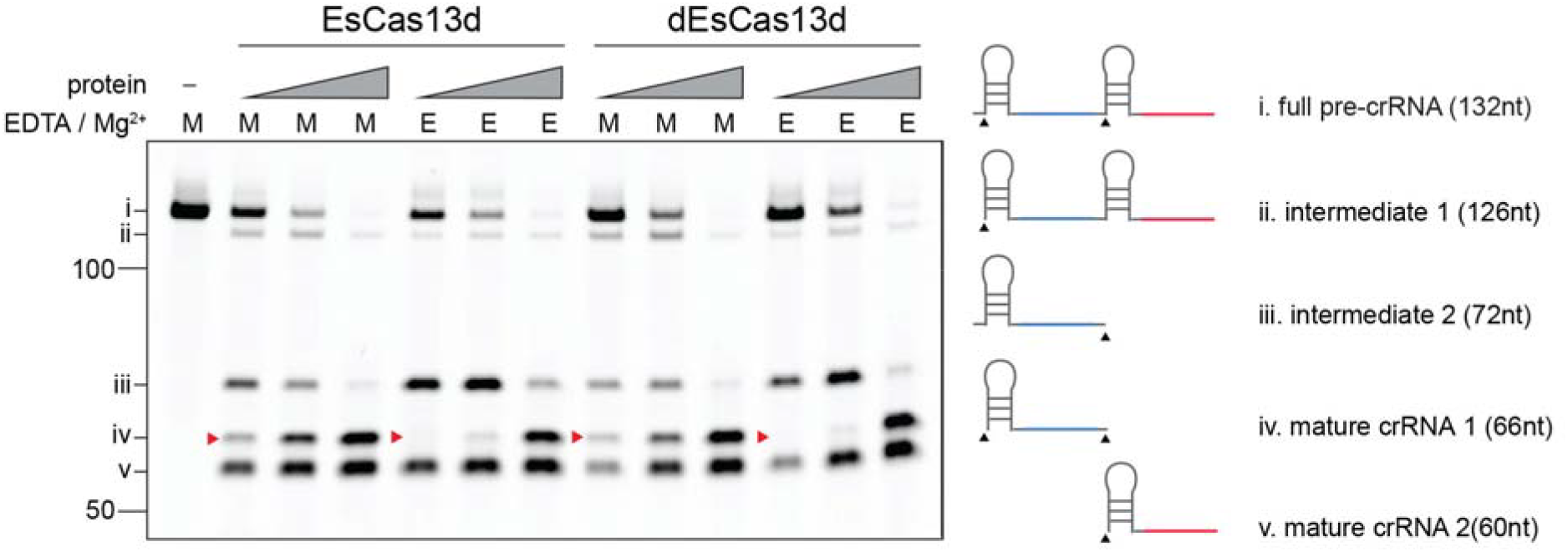
Cas13d pre-crRNA cleavage is enhanced in the presence of Mg^2+^. Denaturing gel of EsCas13d pre-crRNA array cleavage in the presence (M, Mg^2+^) or absence (E, EDTA) of 6 mM Mg^2+^ given an increasing ratio of protein:pre-crRNA (0.5:1, 1:1 and 5:1 molar ratios of EsCas13d to pre-crRNA). Possible cleavage products are indicated as i-v. Catalytic HEPN residues are not required for pre-crRNA processing as d*Es*Cas13d exhibits equivalent activity as active *Es*Cas13d. Red arrowheads indicate the mature crRNA 1 cleavage product requiring two cleavage events. Note the effect of Mg^2+^ at lower protein ratios for this band.

**Figure S5.**
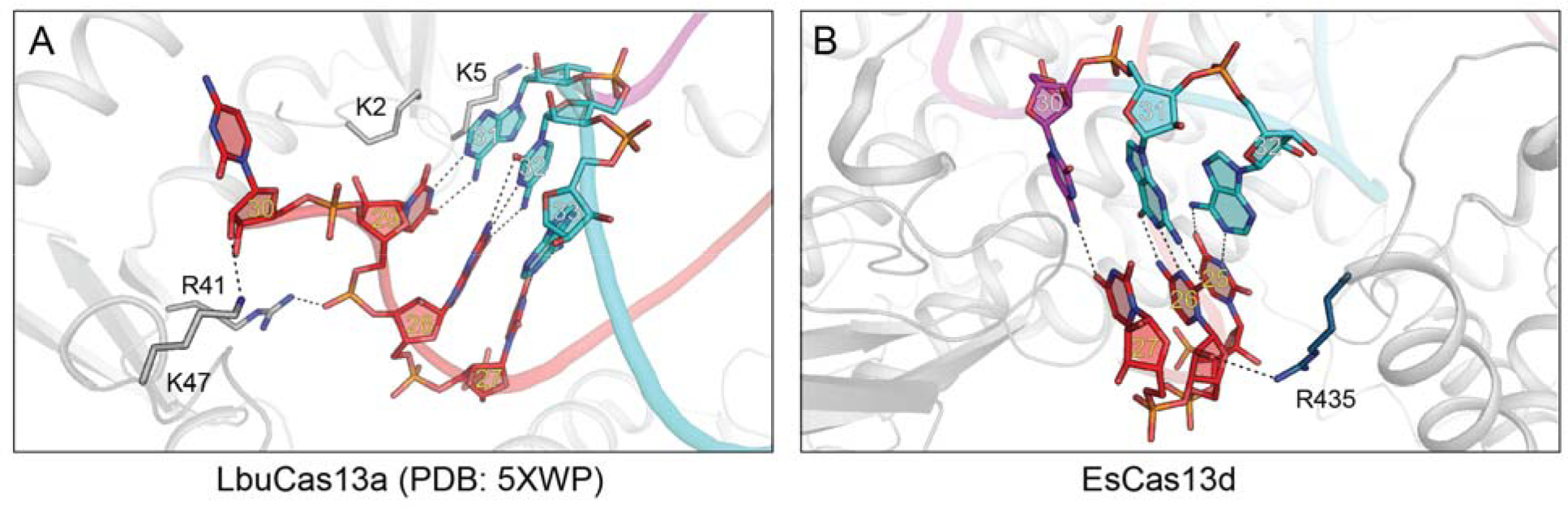
Structural basis for the 3′ PFS requirement in LbuCas13a and EsCas13d. (A-B) Close-up views of 3′ PFS region of LbuCas13a (A) and *Es*Cas13d (B). Key interactions are shown as dashed lines. In *Es*Cas13d, nt-30 faces outward and is free to make base-pairing interactions with incoming target RNA. These would be treated as normal spacer: protospacer interactions or, as in this structure, non-Watson-Crick base-pairs.

**Figure S6.**
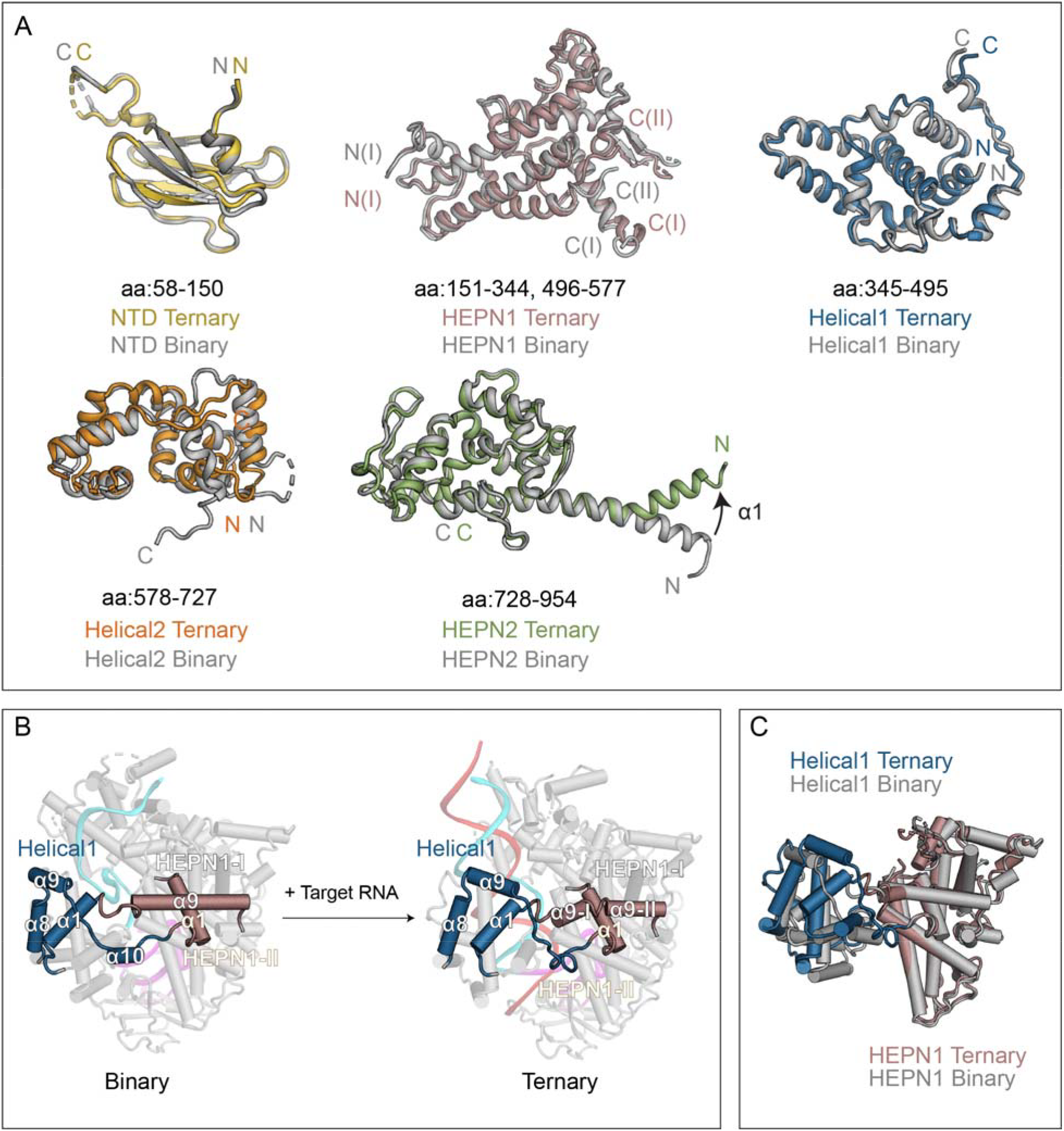
Domain movement of EsCas13d from binary to ternary transition. (A) Superposition of individual *Es*Cas13d domains between binary and ternary forms. (B) Local rearrangement of the hinged region that connects the HEPN1 and Helical-1 domains. (C) Superposition of the Helical-1 and HEPN1 domains between binary and ternary forms. The structures are aligned to HEPN1.

**Figure S7.**
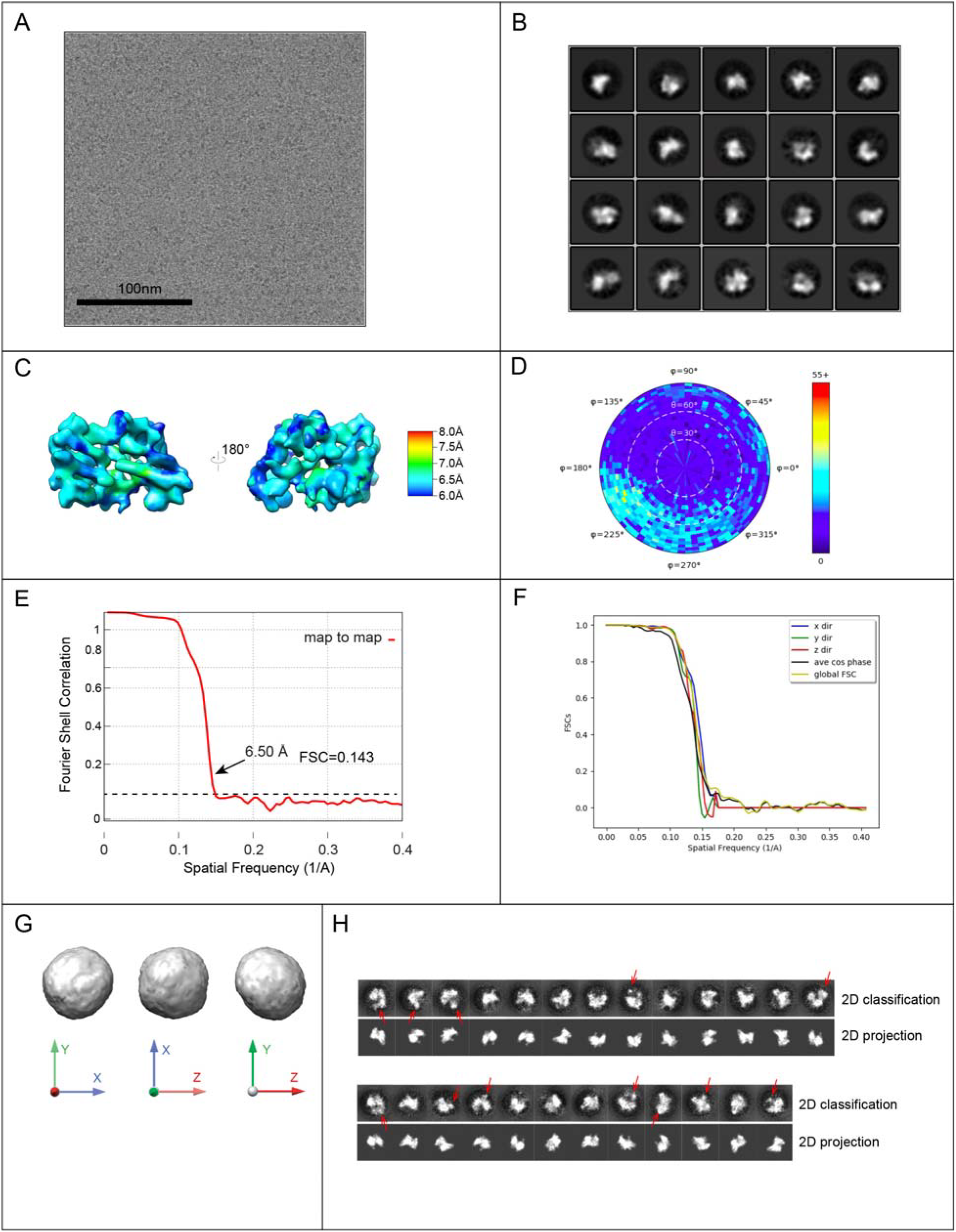
Cryo-EM data collection, analysis, and modeling of Apo *Es*Cas13d. (A) Representative cryo-EM micrograph of Cas13d in the apo form after motion correction. (B) Representative 2D class averages calculated using Relion. (C) Ab initio cryo-EM reconstruction colored by local resolution, calculated using “sxlocres.py” implemented within Sparx. (D) Euler angle distribution plot showing the relative orientation of the particles used in the final 3D reconstruction. (E) Fourier Shell Correlation (FSC) curves calculated between two half maps. (F) 3D FSC curves calculated along z (red), y (green) and x (blue) axes, as well as the global FSC (yellow) and the cosine of the phase residual curves (black) are displayed. (G) 3D FSC isosurfaces, thresholded at a cutoff of 0.75 and displayed in three axial orientations, describe the directional resolution (isotropy) of the refined map. (H) Comparison of *ab initio* 2D class averages calculated using *cis*TEM from the raw data with 2D projections from the apo-*Es*Cas13d model, calculated along optimally assigned Euler angles. Densities that are missing in the 2D projections are often seen within *ab initio* 2D classes averages and are indicated by red arrows.

**Figure S8.**
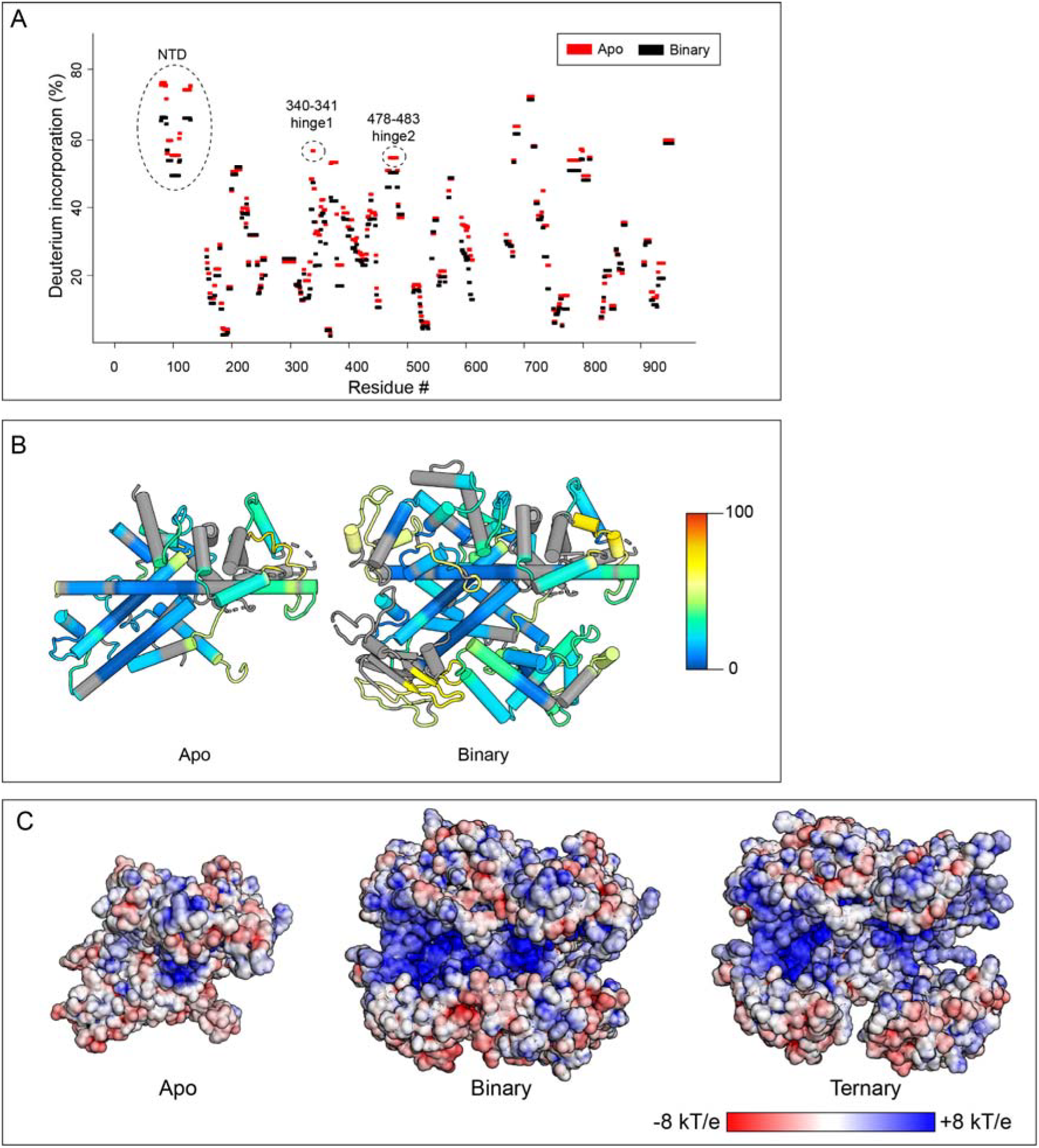
HDX and electrostatic potential maps for *Es*Cas13d. (A) Percentage of deuterium incorporation was consolidated from peptides and plotted as a function of residue number. The amount of deuterium incorporation for each residue, for apo and binary samples, is colored red and black, respectively. Regions with high Deuterium incorporation are indicated. (B) The deuterium uptake for individual apo or binary samples is mapped onto the respective structural models. Blue and red colors represent low and high deuterium uptake. (C) Electrostatic surface maps of *Es*Cas13d in its apo, binary and ternary states. The RNA components are removed for clarity. Red and Blue represent negatively and positively charged regions, respectively.

**Figure S9.**
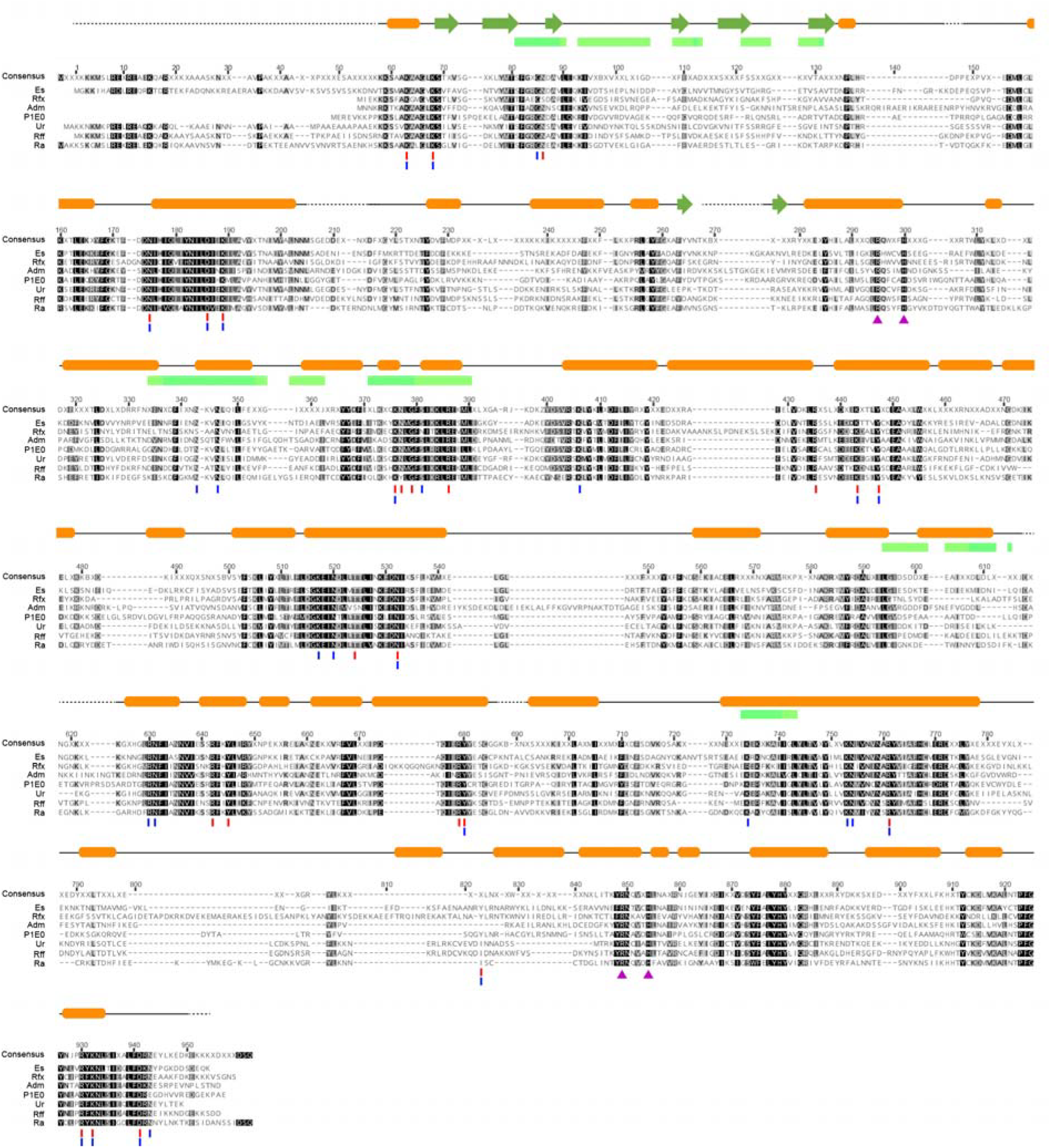
Multiple sequence alignment of Cas13d. Sequence alignment was conducted for seven Cas13d sequences: Es, Rfx, Adm, P1E0, Ur, Rff and Ra. Residue conservation is indicated by grey-scale shading according to Blosum62. Secondary structural elements observed in *Es*Cas13d binary structure are shown above the sequence. Residues that interact with nucleic acids in the binary and ternary states are labeled with blue and red bars, respectively. The same color scheme as Figure 6 was used to highlight the differential HDX onto *Es*Cas13d protein residues.

**Figure S10.**
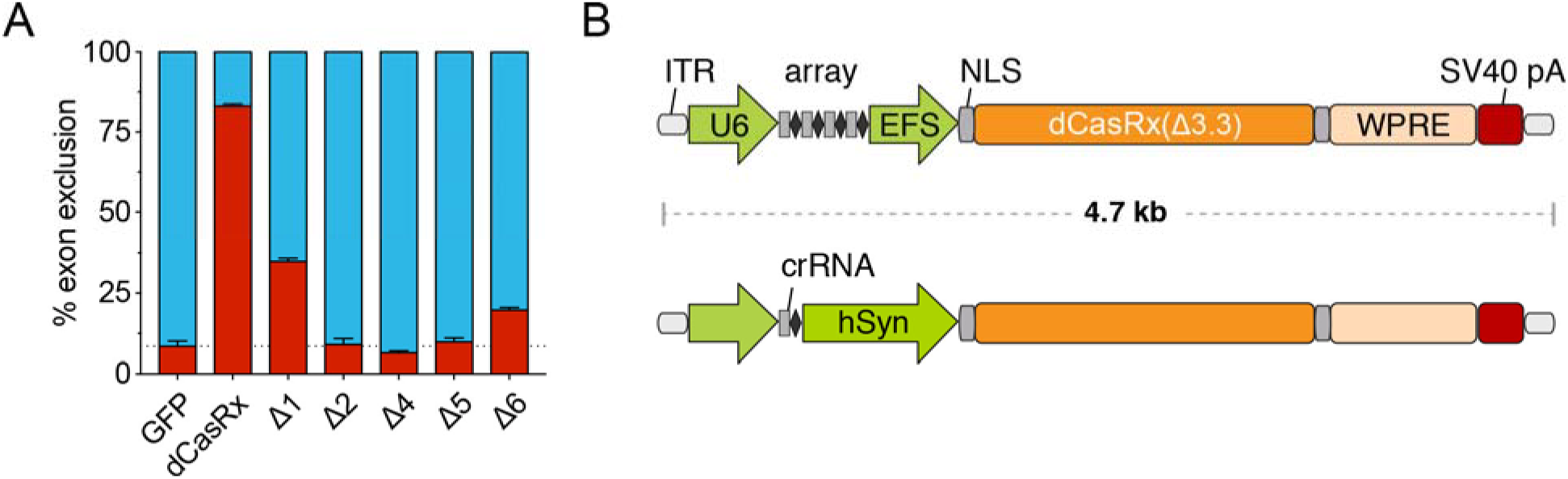
Binding activity and AAV packaging options of truncated CasRx variants. (A) Five CasRx deletion variants from Figure 7B exhibiting minimal cleavage activity were tested on the splicing reporter illustrated in Figure 4F to assay for binding activity. All variants displayed reduced or minimal splicing activity compared to full-length dCasRx. (B) AAV construct designs with Δ3.3 CasRx and a full-length WPRE post-transcriptional element for enhanced transgene expression with payload size <4.7 kb, the AAV packaging limit.

**Supplementary Table 1.** Cryo-EM data collection, image processing, and modeling summary.

| Construct | Cas13d binary complex | Cas13d ternary complex | Cas13d apo form |
| --- | --- | --- | --- |
| **Data collection and Image Processing** |  |  |  |
| Microscope | Arctica Talos | Titan Krios | Arctica Talos |
| Voltage (kV) | 200 | 300 | 200 |
| Camera | Gatan K2 Summit | Gatan K2 Summit | Gatan K2 Summit |
| Defocus range (μm) | 0.8-1.5 | 1.0-2.5 | 1.0-2.0 |
| Defocus mean ± std (μm) | 1.1±0.3 | 1.8±0.4 | 1.5±0.5 |
| Exposure time (s) | 12 | 12 | 12 |
| Dose rate (e<sup>-</sup>/pixel/s) | 2.5 | 3.0 | 2.5 |
| Total dose (e<sup>-</sup>/Å<sup>2</sup>) | 56.8 | 57.3 | 56.8 |
| Pixel size (Å) | 0.73 | 0.79 | 0.73 |
| Exposure rate (ms) | 100 | 200 | 100 |
| Number of micrographs | 1435 | 2205 | 1158 |
| Number of particles (processed) | 412591 | 684169 | 330986 |
| Number of particles (3D classification & refinement) | 207849 | 177910 | 154889 |
| Number of particles (in final map) | 48795 | 51885 | 15846 |
| Symmetry | C1 | C1 | C1 |
| Resolution (global) (Å)* | 3.4 | 3.3 | 6.5 |
| Resolution range (homogeneous portion) (Å) | 3.0-5.0 | 3.0-5.0 | 6.0-8.0 |
| Map sharpening | Spectral flattening<br>between 8Å and 3.4Å | Spectral flattening<br>between 8Å and 3.3Å | Spectral flattening<br>between 10Å and 6.5Å |
| Sphericity @ threshold 0.75 | 0.98 | 0.89 | 0.97 |
| **Model refinement** |  |  |  |
| Number of atoms (modeled) | 8163 | 8834 |  |
| MolProbity score | 1.35 (98<sup>th</sup> percentile) | 1.72 (88<sup>th</sup> percentile) |  |
| Clashscore | 2.85 (98<sup>th</sup> percentile) | 5.78 (91<sup>th</sup> percentile) |  |
| **Protein** |  |  |  |
| Favored rotamers (%) | 98.1 | 94.2 |  |
| Poor rotamers (%) | 0.0 | 0.1 |  |
| Ramachandran favored (%) | 96.0 | 94.0 |  |
| Ramachandran outliers (%) | 0.0 | 0.0 |  |
| Cβ deviations >0.25Å (%) | 0.0 | 0.0 |  |
| Cis Prolines (%) | 0.0 | 0.0 |  |
| Cis Prolines (%) | 0.00 | 0.00 |  |
| Bad bonds (%) | 0.00 | 0.00 |  |
| Bad angles (%) | 0.00 | 0.01 |  |
| **Nucleic acid** |  |  |  |
| Probably wrong sugar puckers (%) | 0.0 | 1.2 |  |
| Bad backbone conformations (%) | 11.76 | 12.05 |  |
| Bad bonds (%) | 0.00 | 0.00 |  |
| Bad angles (%) | 0.00 | 0.03 |  |

\*resolution assessment based on frequency-limited refinement using the 0.143 threshold for resolution analysis

**Movie S1.** Conformational rearrangements between Cas13d binary complex and Cas13d ternary complex, including the binding of target RNA.

## Methods

### Recombinant Cas13d protein purification

EsCas13d and dEsCas13d were purified as previously described (Konermann et al., 2018). In short, Cas13d was expressed as a His-MBP fusion protein with a TEV protease cleavage site in Rosetta2(DE3) cells (Novagen) at 18°C for 20h following IPTG induction. Cells were then lysed in lysis buffer (50 mM HEPES, 500 mM NaCl, 2 mM MgCl2, 20 mM Imidazole, 1% v/v Triton X-100, 1 mM DTT) supplemented with 1X protease inhibitor tablets, 1 mg/mL lysozyme, 2.5U/mL Turbo DNase (Life Technologies), and 2.5U/mL salt active nuclease (Sigma Aldrich) followed by sonication. Following clarification via centrifugation at 18’000g for 1h at 4°C, the lysate was incubated with Ni-NTA Superflow resin (Qiagen) for 1h. The resin was then washed with 10 column volumes of Lysis buffer and eluted with 3 column volumes of elution buffer (50 mM HEPES, 500 mM NaCl, 300 mM Imidazole, 0.01% v/v Triton X-100, 10% glycerol, 1 mM DTT). Samples were dialyzed overnight into into TEV Cleavage Buffer (50 mM Tris-HCl, 250 mM KCl, 7.5% v/v glycerol, 0.2 mM TCEP, 0.8 mM DTT, TEV protease, pH 7.0) before cation exchange with a linear gradient to 1000mM KCl (HiTrap SP, GE Life Sciences) and gel filtration (Superdex 200 16/600, GE Life Sciences). The final purified Protein was frozen for storage at 4 mg/mL in Protein Storage Buffer (50 mM Tris-HCl, 1M NaCl, 10% glycerol, 2 mM DTT, pH 7.5).

### Preparation of guide and target RNAs

*In vitro* transcription template oligos carrying the T7 promoter were synthesized (IDT) and either annealed with an antisense T7 oligo for crRNAs or PCR amplified for targets and arrays. *In vitro* transcription was performed using the Hiscribe T7 High yield RNA synthesis kit (New England Biolabs) at 31°C for 12 hours. crRNAs and short competitors were purified using RNAclean Agencourt AMPure XP beads (Beckman Coulter) with addition of 50% volume of isopropanol for retention of small RNAs. Longer targets and arrays were purified with the MEGAclear Transcription Clean-Up Kit (Thermo Fisher) and frozen at −80°C. The short target for ternary complex formation for cryo-EM imaging was synthesized by Synthego.

*Binary and ternary complex formation* For cryo-EM binary complex formation, 200 μg EsCas13d was incubated with a 3x molar excess of crRNA in complex formation buffer at 37°C for 1 hr (25mM Tris-HCl, 50mM NaCl, 1mM DTT, 1mM MgCl2, pH 7.5). The resulting binary complex was purified by size exclusion chromatography on a Superdex 200 16/600 column (GE Life Sciences) in S200 complex buffer (25mM Tris HCl, 100mM NaCl, 1mM Dtt, MgCl2, 5% glycerol, pH 7.5). Binary peak fractions were pooled and concentrated to 1.5 mg/mL and processed for cryo-EM sample preparation. Ternary formation was performed sequentially, with 15 minutes of binary complex formation at 1:2 ratio of dEsCas13d:crRNA followed by 45 minutes of incubation with target at 1:3 ratio (protein: target). Ternary size exclusion purification and concentration was performed analogous to the binary complex. For HDX binary sample preparation, the purification was scaled up 5X relative to the cryo-EM samples but was otherwise identical. Apo EsCas13d protein was buffer exchanged into S200 complex buffer and normalized to 1.5 mg/mL prior to Cryo-EM sample preparation.

### Electron microscopy sample preparation and data acquisition

All samples, including binary, ternary, and apo, were concentrated to ~1.5mg/ml prior to vitrification. In all 3 cases, Amphipol A8-35 was added to the sample to a final concentration of 0.1%(w/v) immediately before vitrification on cryo-EM grids, in order to ameliorate preferential specimen orientation (Lu et al., 2014). Cryo-EM grids were prepared under >80% humidity at 4°C inside a cold room, and a multi-blotting approach was used to increase particle density (Snijder et al., 2017). Initially, 2ul of sample was applied to an UltrAuFoil R1.2/1.3 300-mesh grid (Quantifoil) after plasma-cleaning (75% argon/25% oxygen atmosphere, 15 W for 7s using a Gatan Solarus). Next, the grid was side-blotted manually with a filter paper (Whatman No.1) followed by a second round of sample loading and side-blot. Finally, another 2ul sample was added to the grid and blotted immediately before plunging into liquid ethane using a manual plunger. Leginon was used for automated EM image acquisition (Suloway et al., 2005). Micrographs of Cas13d-apo and Cas13d binary complex were collected on a Talos Arctica microscope (FEI) operating at 200kV and equipped with a K2 Summit direct electron detector (Gatan). A nominal magnification of 57,000x was used for data collection, providing a pixel size of 0.73 Å at the specimen level, with the defocus range of −0.5 μm to −1.8 μm. Micrographs of Cas13d ternary complex were acquired on a Titan Krios microscope (FEI) operating at 300kV and equipped with a K2 Summit direct electron detector. A nominal magnification of 39,000x was used for data collection, corresponding to a pixel size of 0.79 Å at the specimen level, with the defocus ranging from −1.0 μm to −3.0 μm Movies were recorded in counting mode, with a total dose of ~57e− per Å^2^ for all three samples, fractionated into 120 frames and under a dose rate of ~2.5 – 3 electrons per pixel per second. All details corresponding to individual datasets are summarized in Supplementary Table 1.

### Image processing of Cas13d binary and Cas13d ternary complex

All pre-processing was performed within the Appion suite (Lander et al., 2009). Motion correction was performed using the program MotionCor2 (Zheng et al., 2017) and exposure-filtered in accordance with the relevant radiation damage curves (Grant and Grigorieff, 2015). For processing of Cas13d binary complex and Cas13d ternary complex datasets, structures of LbuCas13a-crRNA complex (PDB:5XWY) and Cas13d-crRNA complex were used as the templates for automatic particle picking in Appion, respectively, using FindEM (Roseman, 2004). The Contrast transfer function (CTF) was estimated using CTFFind4 during data collection on whole micrographs (Rohou and Grigorieff, 2015). After selecting particle coordinates, per-particle CTF estimation was refined using the program GCTF (Zhang, 2016). Stacks containing 400K (Cas13d binary) and 680K (Cas13d ternary) particles were subjected to two rounds of 2D classification, followed by one round of 3D classification in GPU-enabled Relion (Kimanius et al., 2016; Scheres, 2012). The best classes containing 49K (Cas13d binary) and 52K (Cas13d ternary) particles were selected for Relion refinement. Lastly, the parameters were imported into *cis*TEM (Grant et al., 2018), and the last several round of orientation and per-particle CTF refinement were performed to improve the resolution by ~0.1 Å for each dataset.

The spectral amplitudes for each reconstruction were flattened inside *cis*TEM between 8 Å and 3.4 Å or 3.3 Å for the binary and ternary complexes, respectively. The resolutions for both maps were evaluated using conventional Fourier Shell Correlation analysis to evaluate global resolution and directional Fourier Shell Correlation analysis to obtain 3D FSCs and evaluate directional resolution anisotropy (Tan et al., 2017). Due to the manner in which the particles adhered to the air-water interface, the ternary map is characterized by more anisotropic directional resolution.

### Image processing of apo Cas13d

Cryo-EM data was processed in a conceptually similar manner as in binary/ternary. The same templates used for particle picking of Cas13d binary complex were also used to select particles from the apo Cas13d dataset. After CTF estimation in GCTF (Zhang, 2016) and 2 rounds of Relion 2D classification (Scheres, 2012), the extracted stack containing 150K particles was imported into cryoSPARC for *ab initio* reconstruction. We used the following parameters in the reconstruction: Number of Ab-inito classes=1, Initial resolution=20 and Maximum resolution=5, which resulted in a slightly overfitting map with clearly distinguishable secondary structure elements from 16K particles. After a map was generated, the orientations were imported into cisTEM (Grant et al., 2018) without further 3D classification, and the orientations, as well as per-particle CTF parameters were refined for several rounds, resulting in a ~0.2 Å increase in resolution. The final global resolution was estimated at 6.5 Å.

### Model building and refinement

The model of Cas13d binary complex was built *de novo* in Coot (Emsley et al., 2010). A poly-ala model with gaps in looped region was first built based on the EM density, and then residues having bulky side chains (Phe, Trp, and Tyr) were registered to facilitate sequence assignment of the remaining protein. The register of crRNA was conducted based on prior knowledge: that the DR region will form a base-paired stem-loop structure and that the spacer is single stranded. This allowed for unambiguous registration. N-terminal residues 1-57, as well as certain loops scattered throughout the structure and the C-terminal residues 950-954 were poorly ordered, and were thus omitted from the final model. Most of these regions are not conserved among Cas13d orthologs, with ~50% of Cas13d orthologs missing the N-terminal residues (Konermann et al., 2018). For building the model of the Cas13d ternary complex, the binary model was first docked into the ternary cryo-EM map and individual domains were repositioned according to the relevant conformational rearrangements. The NTD, HEPN1, HEPN2 domains and DR of crRNA remain constant, whereas the Helical-1 and Helical-2 domains, as well as the crRNA spacer required repositioning. All connecting loops and any atoms outside of density were rebuilt accordingly. Watson-Crick base pairing between the spacer and target protospacer allowed unambiguous RNA registration. Each model was independently refined in PHENIX (Adams et al., 2010) using phenix.real_space_refine against separate EM half-maps with geometrical, secondary structure, and hydrogen bond restraints. The maps were refined into a working half-map, and improvement of the model was monitored using the free half map. The geometry parameters of the final models were validated in Coot and using MolProbity (Chen et al., 2010). These refinements were performed iteratively until no further improvements were observed. The Cas13d-apo model was generated by rigid-body docking of the Cas13d binary complex structure into the Cas13d-apo cryo-EM map without further refinement or modification, and the parts of the model having poor densities were removed. All the structure figures were prepared in Pymol and UCSF Chimera (Pettersen et al., 2004).

### Biochemical cleavage assays

For DR mutant analysis, purified EsCas13d protein and guide RNA were mixed at 2:1 molar ratio in RNA Cleavage Buffer (25mM Tris pH 7.5, 15mM Tris pH 7.0, 1mM DTT, 6mM MgCl2). Reactions were prepared on ice and incubated at 37°C for 15 minutes prior to the addition of target at 1:2 molar ratio relative to EsCas13d. For pre-crRNA cleavage reactions, purified EsCas13d and EsdCas13d proteins were mixed with purified pre-crRNA at 0.5:1, 1:1 and 5:1 molar ratios in RNA cleavage buffer containing 6mM MgCl2 or EDTA. Reactions were prepared on ice and incubated 37°C for 1 hour. Both cleavage reactions were quenched with 1 μL of enzyme stop solution (10 mg/mL Proteinase K, 4M Urea, 80mM EDTA, 20mM Tris pH 8.0) at 37°C for 15 minutes. The reaction was then denatured with 2X RNA loading buffer (2X: 13mM Ficoll, 8M Urea, 25 mM EDTA), at 85°C for 10 minutes, and separated on a 10% TBE-Urea gel (Life Technologies). DR mutant analysis gels were visualized on the Odyssey Clx Imaging System (Li-Cor); pre-crRNA cleavage gels were stained with SYBR Gold prior to imaging via Gel Doc EZ system (Bio-Rad). For fluorescent ssRNA reporter assays, EsCas13d protein was mixed with guide RNA at 1:1 ratio in RNA cleavage buffer on ice and then assembled into protein:guide complexes at 37°C for 15 minutes. Reactions were put on ice and competitor RNAs were mixed in at 25X molar ratio to target RNA. Target RNAs were added at a 1:5 ratio to EsCas13, and 150nM RNAse-Alert substrate (Thermo-Fisher) was added and mixed. Reactions were then incubated in a real-time PCR machine (Bio-Rad, CFX384 Real-Time System) for 180 minutes at 32 °C and measurements were taken every 5 minutes. Fluorescence values are an average of the last 5 measurements for each condition.

### Cell-based reporter assays

Engineered Cas13d coding sequences were cloned into a standardized expression backbone containing an EF1a promoter and flanking nuclear localization sequences then prepared using the Nucleobond Xtra Midi EF Kit (Machery Nagel) according to the manufacturer’s protocol.

Cas13d arrays were cloned into a minimal backbone containing a U6 promoter. HEK 293FT cells were transfected in 96-well format with 200 ng of Cas13d expression plasmid, 200 ng of guide expression plasmid, and 20 ng of the bichromatic reporter plasmid with Lipofectamine 2000 (Life Technologies). Cells were harvested in FACS Buffer (1X DPBS^−/-^, 0.2% BSA, 2 mM EDTA) after 72 hours, then analyzed in 96-well plate format using a MACSQuant VYB (Miltenyi Biotec) followed by analysis using FlowJo 10. RG6 was a gift from Thomas Cooper (Addgene plasmid # 80167) and modified to replace EGFP with mTagBFP2. All represented samples were assayed with three biological replicates.

